# TPS Proteins coordinate plant growth with sugar availability via the SnRK1 Kinase

**DOI:** 10.1101/2025.08.17.670254

**Authors:** Diana Reis-Barata, Ana Confraria, Leonor Margalha, Bruno Peixoto, Filipa Lopes, Borja Belda-Palazón, Bartimäus Jurke, Stéphanie Arrivault, Vinay Shukla, Regina Feil, Francesco Licausi, Farnusch Kaschani, Markus Kaiser, Camila Caldana, Mark Stitt, John E. Lunn, Elena Baena-González

## Abstract

The ability to sense and respond to nutrients determines adaptation and survival in all organisms. In plants, sucrose stimulates growth and developmental progression via the signalling sugar trehalose 6-phosphate (T6P) which reflects sucrose availability. T6P acts, at least partly, by inhibiting the protein kinase SUCROSE NON-FERMENTING 1 (SNF1)-RELATED KINASE 1 (SnRK1) but the underlying mechanisms are poorly understood. Here, we identify a group of catalytically inactive T6P synthase (TPS) proteins, TPS5/6/7, as important factors for coupling the T6P signal to SnRK1 activity. In *Arabidopsis thaliana,* lack of TPS5/6/7 causes severe growth defects, particularly in roots. This is accompanied by a metabolic signature that is suggestive of T6P insensitivity and impaired sucrose utilization. Using a combination of genetics, SnRK1 activity assays, and imaging, we demonstrate that the growth defects of the *tps5/6/7* mutant are due to SnRK1 misregulation and are reverted by knocking-down SnRK1 in this background. Co-immunoprecipitation assays further show that T6P promotes the interaction of TPS proteins with SnRK1 in a highly specific and dose-dependent manner. Our results support a model where TPS proteins act as T6P sensors, inhibiting non-nuclear SnRK1 activity when sucrose is abundant to promote biosynthetic processes and growth.

## INTRODUCTION

Nutrients sustain life by providing energy and biomass constituents. The ability to sense and respond to nutrient levels determines adaptation and survival and therefore all species have evolved efficient nutrient-sensing mechanisms that suit their needs and environments (Chantranupong et al. 2015). In plants, one crucial nutrient is sucrose, a major product of photosynthesis and the one used by many species to transport carbon from autotrophic source tissues to heterotrophic sink organs (Lunn 2016). Sucrose perception and management are crucial for the proper distribution of carbon and nitrogen resources and are major determinants of stress, growth and developmental responses (Fichtner and Lunn 2021). Changes in sucrose abundance are matched by changes in the signalling sugar trehalose 6-phosphate (T6P) which in turn regulates sucrose levels by preventing further synthesis and by promoting sucrose consumption. Altogether, this maintains sucrose at an optimal level through a system akin to that of insulin and glucose in animals (Yadav et al. 2014; Fichtner and Lunn 2021).

T6P is synthesized from UDP-glucose (UDPG) and glucose 6-phosphate (G6P) by trehalose 6-phosphate synthase (TPS) and is then converted to trehalose by trehalose 6-phosphate phosphatase (TPP) (Cabib and Leloir 1958; Avonce et al. 2006). *Arabidopsis thaliana* (Arabidopsis) has 11 TPS proteins, of which only TPS1 and, to some extent, TPS2 and 4 (referred as class I TPS) have catalytic activity (Vandesteene et al. 2010; Delorge et al. 2015). Class II TPS (TPS5-11, hereafter TPS-II) have an N-terminal TPS-like and C-terminal TPP-like domain. However, they appear to be catalytically inactive (Vogel et al. 2001; Ramon et al. 2009; Vandesteene et al. 2010; Delorge et al. 2015) and have been suggested to play instead signalling functions (Lunn et al. 2014; Van Leene et al. 2022).

The mechanisms by which T6P is sensed and signals downstream are poorly understood but are known to involve the protein kinase SUCROSE NON-FERMENTING 1 (SNF1)-RELATED KINASE 1 (SnRK1) (Zhang et al. 2009; Nunes et al. 2013b; Zhai et al. 2018; Baena-González and Lunn 2020), a conserved key regulator of metabolism and energy homeostasis (Baena-González et al. 2007; Broeckx et al. 2016; Peixoto et al. 2021; Peixoto and Baena-González 2022). SnRK1 is activated under low carbon conditions and is conversely repressed by T6P and other sugar phosphates when carbon levels are high (Baena-González et al. 2007; Zhang et al. 2009; Nunes et al. 2013b; Zhai et al. 2018; Blanford et al. 2024). SnRK1 downregulates growth-related processes through the coordinated control of metabolism and gene expression, playing pivotal roles in energy management during both stress conditions and normal development (Hulsmans et al. 2016; Margalha et al. 2019; Peixoto and Baena-González 2022). Recent work has shown that SnRK1 interacts with TPS-II but the relevance of this interaction remains unclear (Van Leene et al. 2022).

Here, we use a combinatorial knockout approach alongside genetic, biochemical, and plant phenotyping analyses to investigate the function of TPS-II proteins. Plants lacking TPS5/6/7 have aberrantly high T6P and soluble sugar levels, reduced amino acid pools, high oxygen consumption rates and impaired root growth, indicating a disruption in T6P signalling and carbon utilization. We further show that the growth anomalies observed in the *tps5/6/7* mutant result from improper SnRK1 regulation, and that reducing SnRK1 levels in this mutant background restores normal growth. Finally, we demonstrate that T6P promotes the interaction of TPS proteins with SnRK1 in a specific and dose-dependent manner, altogether suggesting that TPS proteins may function as T6P sensors to inhibit non-nuclear SnRK1 activity in accordance with sucrose availability.

## RESULTS

### Lack of TPS5/6/7 impairs root growth

T6P metabolism was recently associated with the regulation of root growth and architecture (Fichtner et al. 2020; Morales-Herrera et al. 2023). This may involve TPS-II proteins, as *TPS-II* promoters are active (Ramon et al. 2009) and TPS-II proteins are present in different root regions (Mergner et al. 2020). Of all TPS-II proteins, TPS7 is the most abundant one across different Arabidopsis tissues, including whole roots and root tips (Mergner et al. 2020)(Supplementary Fig. 1). We therefore initially focused on TPS7 and compared root growth in control plants (Col-0) and a *tps7* loss-of-function mutant (Fig. 1A and Supplementary Fig. 2). To minimize the potential impact of varying germination times on root size, seedlings were grown vertically on 0.5× MS plates for 7 d and seedlings with comparable root size were thereafter transferred to new plates and grown for an additional 7 days. Primary root (PR) growth was measured as the length of root developed post-transfer. The *tps7* mutant displayed a small but significant reduction in PR length compared to the Col-0 control and this phenotype was rescued in two independent complementation lines expressing mCherry-tagged TPS7 under the control of the *TPS7* promoter and terminator (*TPS7-mCherry, C#1* and *C#2*, Fig. 1A).

**Figure 1.**
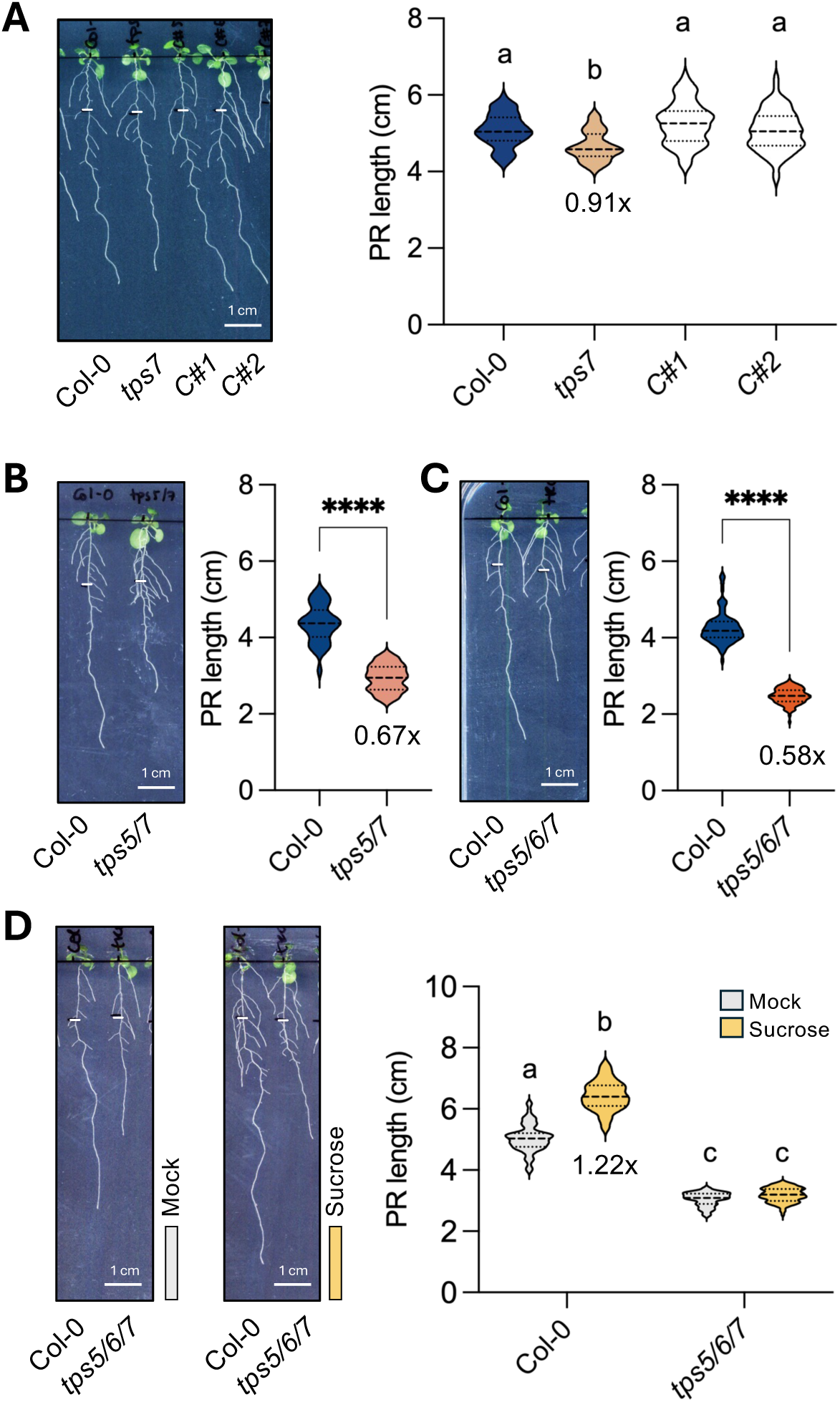
TPS5/6/7 are important for root growth. Representative images of 14d-old seedlings grown vertically on 0.5× MS medium. (**A**) Col-0, *tps7* and *proTPS7::TPS7-mCherry* (*C#1* and *C#2*), (**B**) Col-0 and *tps5/7*, (**C,D**) Col-0 and *tps5/6/7*. After 7d of growth seedlings were transferred to fresh 0.5× MS medium supplemented (**D** middle) or not (**A-D** left) with 30 mM sucrose. White marks indicate the position of the root tip at transfer. Scale bar 1 cm. Right (**A-D**), quantification of primary root (PR) length, measured from the white mark to the root tip. Number of independent experiments and seedlings per genotype: (**A**) *n*=3, 58-67 seedlings; (**B**) *n*=2, 34 seedlings; (**C**) *n*=6, 79-100 seedlings; (**D**) *n*=3, 76-85 seedlings. The dotted lines represent the quartiles and the dashed line, the median. Different letters (**A,D**) or asterisks (**B,C**) indicate statistically significant differences between genotypes (*p*<0.05, Brown-Forsythe and Welch ANOVA with Dunnett’s T3 test in **A**, Kruskal-Wallis with Dunn’s test in **D**; two-tailed unpaired Student t-test in **B** and Welch’s t-test in **C**, *p<0.05, **p<0.01, ***p<0.001).

Given that TPS5 and TPS6 are phylogenetically closest to TPS7 (Avonce et al. 2006; Lunn 2007), we next investigated whether they also contribute to root growth. Individual *tps5* and *tps6* loss-of-function mutants (Supplementary Fig. 2) did not show statistically significant differences in PR length compared to Col-0, although a slight growth reduction was observed for *tps5* (Supplementary Fig. 3). To assess potential functional redundancy, we also examined double *tps5/7* and triple *tps5/6/7* mutants. The phenotypes were more pronounced in higher-order mutants and increased in severity with the progressive accumulation of *tps* mutations, with PR length in the *tps5/6/7* mutant being only 58% of that in Col-0 (Fig. 1B-C). Although at this developmental stage, no obvious impact was observed in the growth of the aerial part, a smaller rosette size was evident at a later stage when plants were grown on soil (Supplementary Fig. 4).

Given the involvement of T6P metabolism in carbon allocation, we reasoned that the stunted root phenotype of the *tps5/6/7* mutant could result from disrupted sugar provision from source leaves to roots. To test this, Col-0 and *tps5/6/7* seedlings were grown vertically on 0.5× MS plates supplemented or not with 1% sucrose. As expected, in Col-0 seedlings, the presence of sucrose induced a significant increase in PR length compared to mock plates. However, the PR length of the *tps5/6/7* mutant remained unaffected by sucrose (Fig. 1D). In heterotrophic tissues, the catabolism of sucrose provides crucial substrates for primary metabolism, including glucose and fructose, thus supporting cellular maintenance and growth. We therefore next investigated whether the growth defects of the *tps5/6/7* mutant could be rescued by supplementing its breakdown products. However, the addition of glucose and fructose promoted PR growth in Col-0 but it had no effect in the *tps5/6/7* mutant (Supplementary Fig. 5).

Altogether, our results show that the stunted root growth of the *tps5/6/7* mutant is not due to impaired sugar provision to this organ but is rather caused by an inability to respond to sugar availability.

### Lack of TPS5/6/7 causes aberrantly high sucrose, T6P, and T6P:sucrose ratios

To investigate the cause for the impaired growth of the *tps5/6/7* mutant and its lack of response to sugar, we first performed metabolite analyses. To avoid sugar carry-over from the medium, we tested whether growing seedlings in mock plates but under contrasting light intensities (Control light, 110 μmol m^−2^ s^−1^; High Light, 200 μmol m^−2^ s^−1^) could reproduce the differences in PR growth observed between genotypes in the absence or presence of exogenous sugars. Growth under high light intensity led to a significant increase in PR length in Col-0 but had no effect in the *tps5/6/7* mutant (Fig. 2A), hence recapitulating, albeit to a milder extent, the effect of sugar supplementation (Fig. 1D). We then grew seedlings under both control and high light conditions and harvested samples at the end of the night (EN), when growth rates are known to increase strongly in roots (Yazdanbakhsh et al. 2011) and shoots (Nozue et al. 2007). To minimize the masking of potential tissue-specific responses, we collected shoots, roots, and root tips (<0.5 cm) separately and analysed metabolites related to carbon and nitrogen primary metabolism by LC-MS/MS.

**Figure 2.**
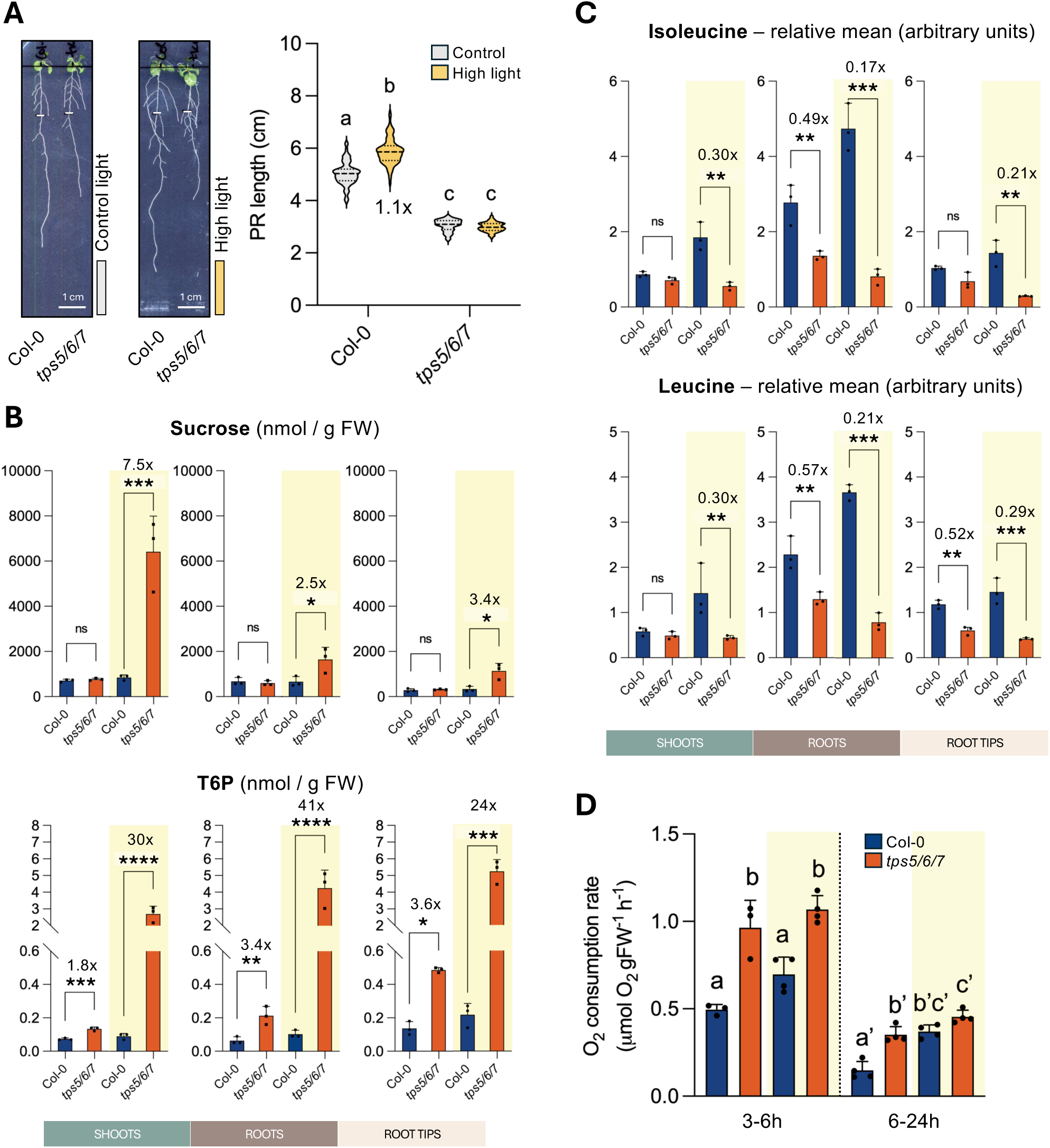
Lack of TPS5/6/7 causes a starvation-like metabolic syndrome. (**A**) Left and middle, representative images of 14d-old Col-0 and *tps5/6/7* seedlings grown vertically on 0.5x MS medium under control light conditions (110 μmol m^−2^ s^−1^) for 7 d, transferred to fresh 0.5x MS plates and grown for 7 d under control light (CL; left) or high light conditions (HL; 200 μmol m⁻² s⁻¹; right). Scale bar 1 cm. Right, quantification of PR length, measured from the white mark to the root tip (*n*=3 independent experiments, 76-82 seedlings per genotype per condition). The dotted lines represent the quartiles and the dashed line, the median. Different letters indicate statistically significant differences between genotype and condition (*p*<0.05, Kruskal-Wallis with Dunn’s test). (**B-C**), Levels of sucrose and T6P (**B**), and Isoleucine and Leucine (**C**) in the indicated tissues of 14d-old Col-0 and *tps5/6/7* seedlings grown as in (A) under contrasting light intensities (CL, white; HL, yellow). Graphs show the mean of 3 independent experiments (each consisting of 200-250 seedlings per genotype per condition, error bars, SD). Asterisks indicate statistically significant differences between genotypes within each tissue and condition (two-tailed unpaired Student and Welch’s t-tests, *\*p*<0.05, *\*\*p*<0.01, *\*\*\*p*<0.001). (**D**) Oxygen consumption rates (μmol O₂ g⁻¹ FW h⁻¹) of 10d-old Col-0 and tps5/6/7 seedlings upon transfer to fresh 0.5x MS medium with (yellow) or without sucrose (white). Shown are rates in the 3-6h and 6-24h time windows. Bars show the mean of 4 independent experiments (error bars, SD). Different letters indicate statistically significant differences between genotype and condition (*p*<0.05, one-way ANOVA with Tukey HSD test).

In Col-0, sucrose levels were lowest in root tips while T6P levels were highest in this tissue (Fig. 2B). This resulted in higher T6P:sucrose ratios in root tips (Table 1) than in other tissues, consistent with the higher T6P:sucrose ratios reported in actively growing sinks (Lunn et al. 2014). Under control light, sucrose levels were similar in Col-0 and *tps5/6/7* but T6P levels were significantly higher in the mutant in all tissues (Fig. 2B). Under high light, sucrose levels were higher in all *tps5/6/7* tissues (Fig. 2B) and this was accompanied by a greater accumulation of the sucrose synthesis intermediate, sucrose 6-phosphate (Supplementary Fig. 6). Under high light the differences between genotypes were dramatically larger for T6P, mounting up to 41-fold in roots (Fig. 2B). This resulted in significantly higher T6P:sucrose ratios in the *tps5/6/7* mutant across all tissues and light conditions (Table 1). In all Col-0 tissues, both sucrose and T6P levels were in a similar range in the control and high light samples, as expected from the tight coupling between the two sugars (*ns*, Student t-test). However, this was not the case in the mutant where sucrose and T6P were significantly higher in the high light than the control light samples across all tissues (*p*<0.05, Student t-test).

**Table 1.**
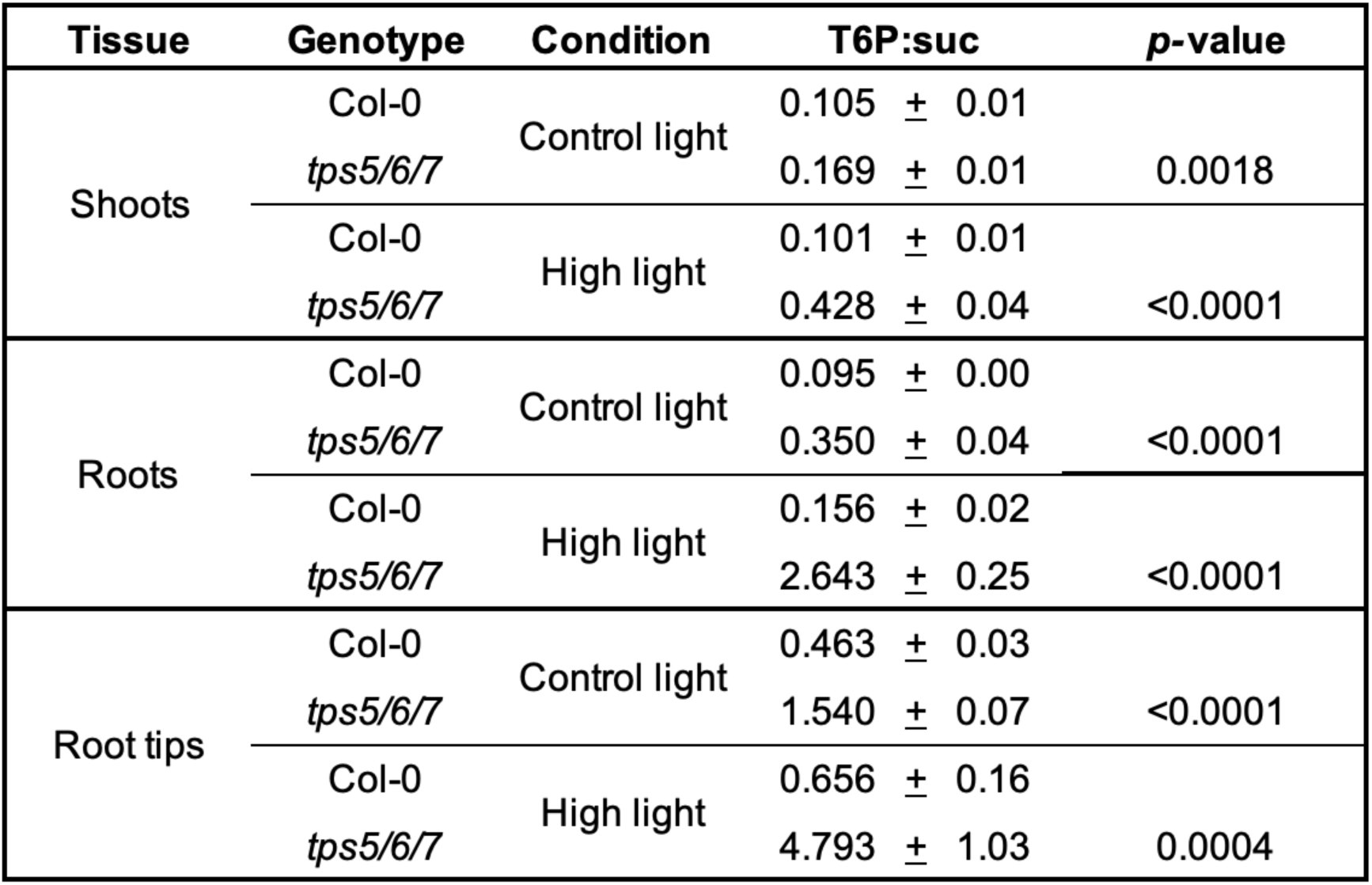
Impact of TPS5/6/7 on T6P:sucrose ratios. T6P:sucrose ratios are means ± SD and were calculated using the T6P and sucrose values represented in Fig. 2B. *p-*values refer to statistically significant differences between Col-0 and *tps5/6/7* for each growth condition within the different tissues (two-tailed unpaired Student t-test).

The *tps5/6/7* mutant also had higher levels of the sucrose degradation products, glucose and fructose (Supplementary Fig. 6), consistent with the inability of these sugars to rescue the growth defects of the mutant (Supplementary Fig. 5). Under control light, glucose and fructose were moderately elevated in *tps5/6/7* root tissues, but the differences were markedly exacerbated under high light (Supplementary Fig. 6). Aberrantly high levels were also observed for trehalose, the degradation product of T6P, whose pattern closely followed that of T6P (Supplementary Fig. 6).

Collectively, our results reveal substantially altered sucrose and T6P metabolism in the *tps5/6/7* mutant, leading to dramatically increased T6P:sucrose ratios, particularly under high light conditions. This, together with the elevated levels of sucrose and T6P breakdown products and the inability of the *tps5/6/7* mutant to promote root growth, suggests a potential disruption in T6P signalling and sugar utilization.

### Lack of TPS5/6/7 causes a metabolic syndrome suggestive of altered T6P signalling

Previous studies in Arabidopsis have reported that an inducible rise in T6P leads to major metabolic changes, including increased accumulation of organic and amino acids (Figueroa et al. 2016; Ishihara et al. 2022; Avidan et al. 2024). Consequently, we investigated whether the elevated T6P levels observed in the *tps5/6/7* mutant (Fig. 2B) elicited similar metabolic alterations in our samples.

Despite the substantial accumulation of T6P and sugars in the mutant, most central metabolites remained similar between Col-0 and *tps5/6/7* (Supplementary Fig. 7A; Supplementary Tables S1 and S2). Within glycolysis, significant differences were observed only for fructose 1,6-bisphosphate (Fru1,6BP) and pyruvate which were at higher levels in *tps5/6/7 vs*. Col-0 roots under high light and for glucose 6-phosphate (G6P) and phospho*enol*pyruvate (PEP) whose levels were lower in *tps5/6/7 vs*. Col-0 shoots under control light. For TCA cycle intermediates the differences were moderate and varied across tissues. In shoot samples, the *tps5/6/7* mutant had significantly higher levels of most organic acids than Col-0 under high light (Supplementary Fig. 7A). In root tips, on the other hand, the levels were mostly lower in the mutant, with significant decreases occurring for malate, oxoglutarate (2-OG), and fumarate under control light and for malate and aconitate under high light.

To gain further insight into the metabolic alterations of the *tps5/6/7* mutant, we used GC-MS/MS to measure the relative levels of amino acids. Most amino acids were at lower levels in the mutant tissues, especially under high light (Supplementary Fig. 7A), with particularly pronounced decreases for lysine (Lys) and branched-chain amino acids [BCAAs, leucine (Leu), isoleucine (Iso) and valine (Val)] (Fig. 2C; Supplementary Fig. 7B). Under control conditions, the levels of Leu and Iso in mutant roots were roughly half of those in Col-0. Under high light, these differences were strongly accentuated, becoming evident in all tissues (ranging from 30% to 17% of Col-0 levels) and affecting also Val and Lys (Fig. 2C; Supplementary Fig. 7B). In contrast, three amino acids exhibited significant changes in the opposite direction under high light, with proline (Pro) being significantly elevated in shoots and root tips, and alanine (Ala) and glutamine (Gln) in roots in *tps5/6/7* compared to Col-0 (Supplementary Fig. 7). Despite these changes in free amino acids, total protein levels appeared similar between Col-0 and *tps5/6/7* across all tissues and conditions (Supplementary Fig. 8).

The degradation of sugars and amino acids generates TCA cycle intermediates that can be used for ATP production in mitochondrial respiration. We therefore asked whether the accumulation of sugars in the *tps5/6/7* mutant and its reduced growth could be related to altered respiration. To this end, we measured oxygen consumption rates in 10d-old seedlings grown with or without 1% sucrose. Measurements were taken at ZT4, immediately after transferring seedlings to an air-tight vial sealed with a septum for oxygen electrode insertion, and subsequently at 3 h, 6 h and 24 h post-transfer. Oxygen consumption rates were calculated for the 3-6 h and 6-24 h intervals (Fig. 2D). In the 3-6 h interval, the rate of oxygen consumption was significantly higher in *tps5/6/7* seedlings (Fig. 2D). This was true both in the absence and presence of exogenous sucrose (1.94-fold and 1.53-fold increases, respectively), suggesting that the differences are not due solely to the higher endogenous sugar levels of the mutant. In the 6-24 h interval, overall rates were lower (>50%) than in the 3-6 h interval, possibly reflecting the reduced metabolic demands of plants under prolonged darkness. In the absence of sucrose, rates were significantly higher in *tps5/6/7* seedlings (2.4-fold), but in the presence of sucrose the rates were similar between both genotypes. As expected, sucrose stimulated oxygen consumption in Col-0 (1.4-fold at 6 h and 2.5-fold at 24 h). However, the effect of sucrose was clearly weaker in *tps5/6/7* seedlings (1.1-fold at 6 h and 1.3-fold at 24 h), with the difference between Col-0 and *tps5/6/7* reaching significance in the 6-24 h interval (*p*=0.03, Student *t*-test).

Altogether, the metabolic profile of the *tps5/6/7* mutant does not align with expectations based on its elevated sucrose and T6P levels, suggesting that lack of TPS-II proteins causes T6P insensitivity and impaired sucrose utilization.

### Lack of TPS proteins alters SnRK1 function

The SnRK1 protein kinase is known to be induced by carbon starvation (Baena-González et al. 2007; Mair et al. 2015; Pedrotti et al. 2018; Ramon et al. 2019). Furthermore, TPS proteins were shown to interact with various subunits of the SnRK1 complex (Van Leene et al. 2022). We therefore next investigated whether the phenotypes of the *tps5/6/7* mutant were related to SnRK1 function. To this end, we crossed *tps5/6/7* plants to a partial loss-of-function mutant of the SnRK1α catalytic subunit, which is encoded by two genes (*SnRK1α1* and *SnRK1α2*) in Arabidopsis. From the cross with the SnRK1 *sesquiα* mutant (*snrk1α1*^-/-^; *snrk1α2*^-/+^), we retrieved *tps5/6/7/snrk1α1* and *tps5/6/7/sesquiα* progeny which were used alongside the corresponding parental lines in PR phenotyping assays. The depletion of SnRK1α function markedly alleviated the PR growth defects of the *tps5/6/7* mutant, with the *snrk1α1* and *sesquiα* mutations restoring the *tps5/6/7* root growth defects from 58% to 72% and 92%, respectively, of the Col-0 levels (Fig. 3A). This shows that the growth defects of the *tps5/6/7* mutant are largely SnRK1-dependent and suggests TPS5/6/7 are negative regulators of SnRK1 function.

**Figure 3.**
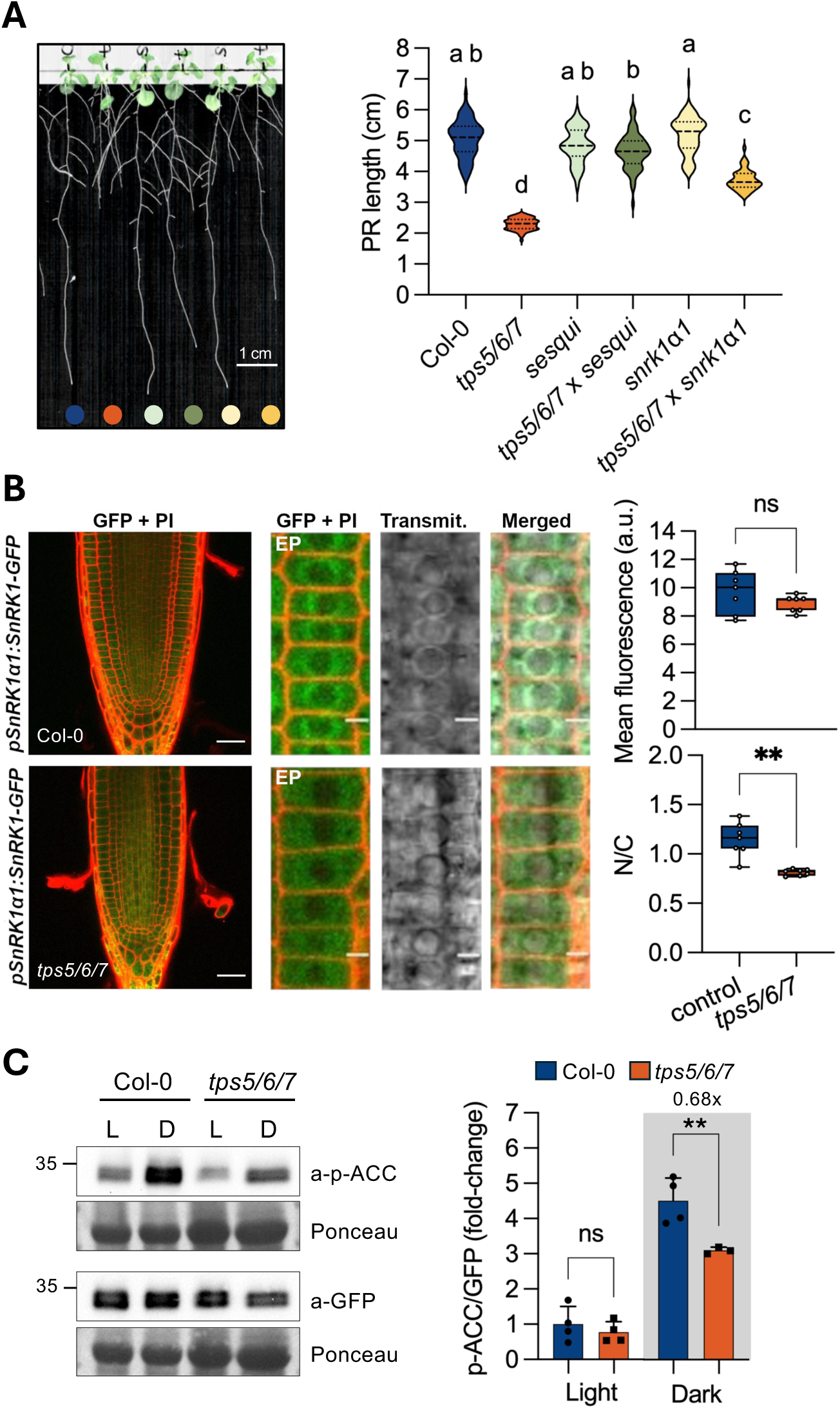
Lack of TPS proteins alters SnRK1ɑ1 function. (**A**) *Left*, representative images of 14-d old Col-0, *tps5/6/7*, *sesqui*, *tps5/6/7 x sesqui*, *snrk1ɑ1* and *tps5/6/7 x snrk1ɑ1* seedlings grown vertically on 0.5x MS medium for 7 d and transferred to fresh 0.5x MS plates for 7 d. Scale bar 1 cm. *Right*, quantification of PR length, measured from the white mark to the root tip (n=3; 54-92 seedlings per genotype). The dotted lines represent the quartiles and the dashed line, the median. Different letters indicate statistically significant differences between genotypes (*p*<0.05, Kruskal-Wallis with Dunn’s test). (**B**) Analyses of SnRK1α1 subcellular localization by CLSM in root apical meristems of 8d-old SnRK1α1-GFP and *tps5/6/7* SnRK1α1-GFP seedlings stained with propidium iodide (PI). Left, representative images of SnRK1α1 localization in the root apex. Right, quantification of mean cellular fluorescence (up) and nucleus-to-cytosol ratio (N/C) ratio (down) of SnRK1α1-GFP in the indicated genotypes (*n*=7 root tips, each consisting of the average of 5 meristematic cells). Lower and upper box boundaries represent the first and third quantiles, respectively, horizontal lines mark the median and whiskers mark the highest and lowest values. *P*-values denote statistically significant differences (two-tailed unpaired Welch’s t-test). EP, epidermis; a.u., arbitrary units. Scale bars, 30 µm or 5 µm for photographs of the whole root apex or epidermal cells (EP), respectively. (**C**) Left, representative immunoblots showing in vivo phosphorylation of a nuclear rat ACC-GFP-HA reporter protein in 14d-old Col-0 and *tps5/6/7* seedlings, either maintained in the light (L) or transferred to darkness (D) at ZT2 for 4 h and harvested at ZT6. GFP immunodetection was used for normalization. Right, quantification of SnRK1ɑ1 nuclear activity, presented as the p-ACC/GFP ratio. Bars show the mean of 4 independent experiments (error bars, SD). Asterisks indicate significant differences between genotypes (*\*p*<0.05, *\*\*p*<0.01, *\*\*\*p*<0.001; two-tailed unpaired Student t-test).

To investigate which aspects of SnRK1 activity are altered in the absence of TPS5/6/7, we first compared total SnRK1α1 levels and SnRK1α T-loop phosphorylation, which is essential for SnRK1 activity (Baena-González et al. 2007; Shen et al. 2009). This revealed no differences between Col-0 and the mutant (Supplementary Fig. 9). We next compared the subcellular localization of SnRK1α1 in Col-0 and *tps5/6/7* seedlings, given that SnRK1 function is largely determined by its subcellular localization (Ramon et al. 2019; Gutierrez-Beltran and Crespo 2022). To this end, we crossed the *tps5/6/7* mutant with a line expressing SnRK1α1-GFP under the control of *SnRK1α1* upstream and downstream regulatory regions (Crozet et al. 2016) and imaged roots of the fully homozygous progeny at the end of the night using confocal laser scanning microscopy (CLSM). The total mean fluorescence was similar in the two genotypes (Fig. 3B, upper panel), confirming the immunoblot analyses (Supplementary Fig. 9). As previously reported (Ramon et al. 2019; Belda-Palazón et al. 2020, 2022), in Col-0 roots, SnRK1α1 was present in the cytoplasm and was strongly enriched in the nucleus. In the *tps5/6/7* mutant, however, the nucleus-to-cytosolic ratio was 30% lower than in Col-0 (Fig. 3B, lower panel), pointing to a redistribution of SnRK1α1 from the nucleus to the cytosol. To compare SnRK1α1 localization to that of TPS7, we next imaged our *TPS7-mCherry* complementation line *C#2* (Fig. 1A) under the same conditions as SnRK1α1-GFP. This revealed a fully extranuclear localization for TPS7 (Supplementary Fig. 10).

To assess the possible functional consequences of altered SnRK1α1 localization, we crossed the *tps5/6/7* mutant with a reporter line expressing a peptide from rat acetyl-CoA carboxylase (ACC) which is specifically phosphorylated by SnRK1 (Muralidhara et al. 2021; Sanagi et al. 2021). The reporter protein is localized to the nucleus and therefore its relative phosphorylation serves as readout for SnRK1 nuclear activity (Muralidhara et al. 2021). We subjected *NLS-ACC* and *tps5/6/7/NLS-ACC* seedlings to an “energy stress” treatment (4h of sudden darkness) which triggers nuclear SnRK1 signalling (Margalha et al. 2023). The treatment increased NLS-ACC phosphorylation in both lines, indicating the induction of SnRK1 nuclear activity (Fig. 3C). However, the phosphorylation of the reporter under energy stress conditions was significantly lower in the *tps5/6/7* mutant. This indicates that SnRK1 nuclear activity is weaker in *tps5/6/7* seedlings and is consistent with the reduced nuclear localization of SnRK1α1 in the mutant (Fig. 3B).

### T6P enhances the interaction of TPS proteins with SnRK1

TPS proteins have a phosphatase-like domain that retains key residues for T6P binding (Lunn 2007). Given their physical interaction with SnRK1 (Van Leene et al. 2022), the insensitivity of the *tps5/6/7* mutant to T6P (Fig. 2) and the inhibition of SnRK1 by T6P (Zhang et al. 2009; Nunes et al. 2013b; Zhai et al. 2018; Baena-González and Lunn 2020), we next wondered whether the interaction of TPS proteins with SnRK1 is affected by T6P. To this end, we performed co-immunoprecipitation experiments using our *TPS7-mCherry* complementation lines (Fig. 1A). We grew *TPS7-mCherry* (C#2) seedlings in mock plates, immunoprecipitated TPS7 in the absence or presence of T6P, and assessed the interaction with endogenous SnRK1α1 by immunoblotting (Fig. 4A and Supplementary Fig. 11A). The presence of T6P increased in a clear dose-dependent manner the interaction between TPS7 and SnRK1α1. To further assess the specificity of this effect, we next performed similar experiments in the presence of other sugars linked to T6P metabolism (sucrose, UDPG, G6P and Trehalose) or previously shown to inhibit SnRK1 [G6P and G1P; (Nunes et al. 2013b)]. However, the interaction between SnRK1 and TPS7 was not affected by any of the other tested sugars, indicating that the effect of T6P is highly specific (Fig. 4B). To further confirm the effect of T6P in the TPS7-SnRK1 interaction we performed reciprocal co-immunoprecipitation analyses using a SnRK1α1-GFP complementation line (Crozet et al. 2016) followed by tandem mass spectrometry (MS/MS) analyses of the SnRK1α1-GFP protein interactors (Supplementary Fig. 11B). The presence of T6P significantly increased the interaction of SnRK1α1-GFP with TPS7 but also with other TPS proteins, corroborating the results obtained with the *TPS7-mCherry* line and suggesting a broader role for T6P in regulating the interaction of SnRK1 with the whole TPS-II protein family (Fig. 4C).

**Figure 4.**
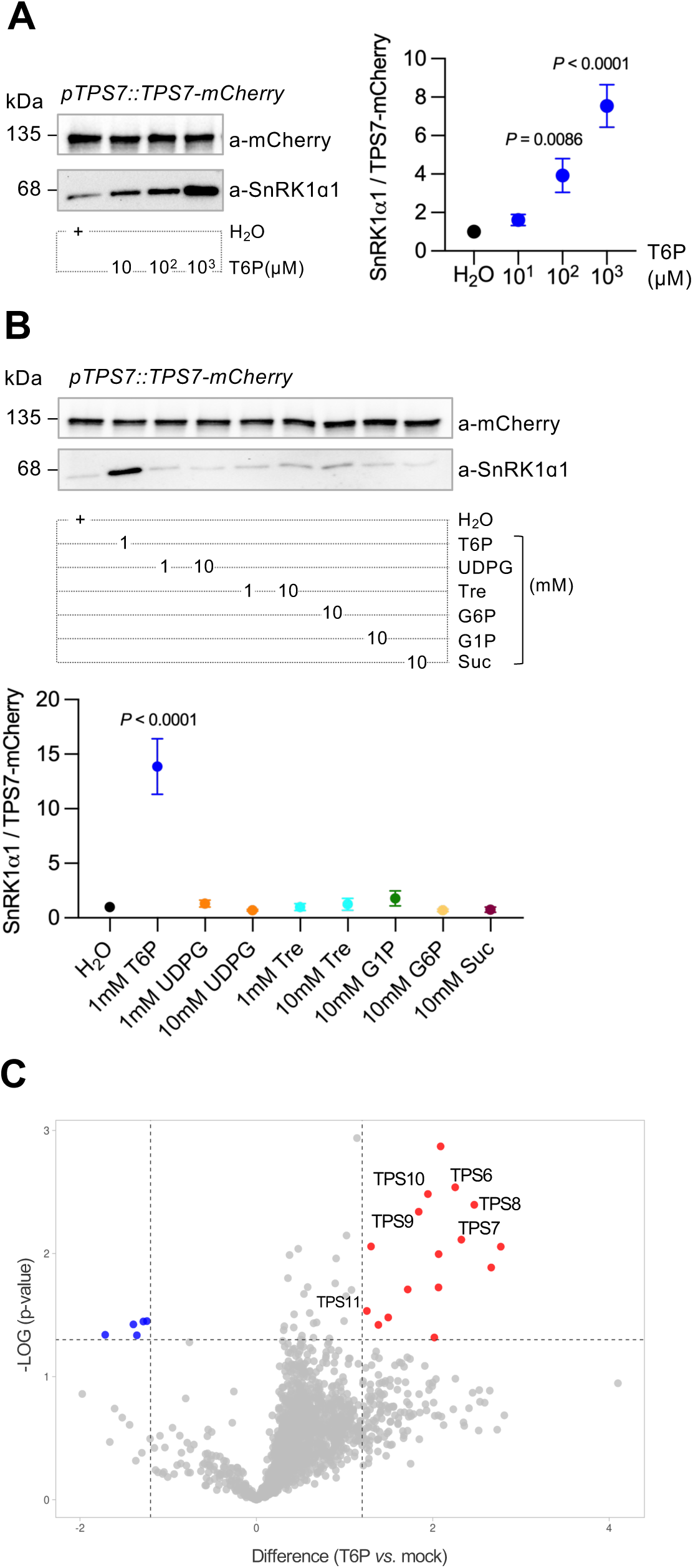
T6P enhances TPS-SnRK1α1 interactions. (**A**,**B**) Up, representative western blots (WB) of immunoprecipitated TPS7-mCherry (top) and co-immunoprecipitated SnRK1ɑ1 (bottom) in the presence of the indicated concentrations of T6P (**A**) and/or other sugars (**B**) (experimental set-up in Supplementary Fig. 11A). Bottom, relative quantification of SnRK1ɑ1 amounts co-immunoprecipitated with TPS7-mCherry in the indicated conditions. Graphs show the mean of 3 independent experiments (error bars, SEM). *P* values denote statistically significant differences between control and treated samples (one-way ANOVA with Dunnet’s test). (**C**) Volcano plot showing changes in SnRK1ɑ1 interactors in the presence of T6P as per the experimental setup in Supplementary Fig. 11B. The x-axis represents the Difference (LFQ T6P - LFQ mock), and the y-axis indicates the −log10(*P*-value). Points represent individual interactors, with the ones significantly changed highlighted in blue (depleted) or red (enriched).

Altogether, our results show that T6P enhances the TPS-SnRK1 interaction, suggesting that TPS proteins may serve as T6P sensors that inhibit SnRK1 activity.

## DISCUSSION

Here, we demonstrate that Arabidopsis TPS-II proteins TPS5/6/7 play an important role in T6P signalling, coordinating metabolism and growth with sucrose supply through inhibition of the SnRK1 kinase (Fig. 5).

**Figure 5.**
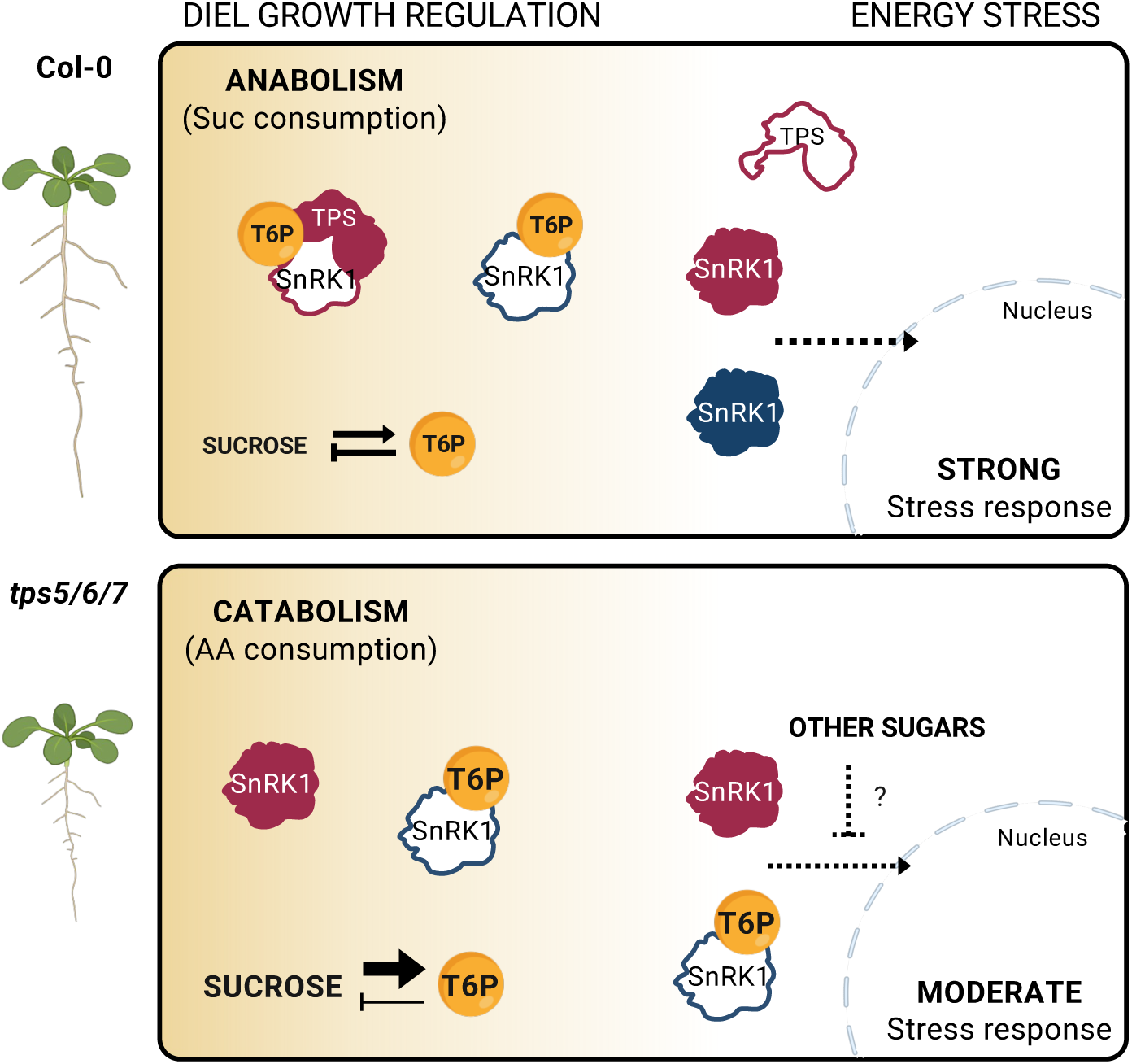
TPS-II proteins coordinate plant growth with sugar availability via the SnRK1 kinase – a model. Up left, in wild-type plants (Col-0) growing under favourable conditions, high T6P enhances the TPS-SnRK1 interaction, leading to inhibition of a non-nuclear SnRK1 pool. Other (non-nuclear) pools of SnRK1 may be directly inhibited by T6P or other signals/factors. Low SnRK1 activity promotes sucrose consumption through anabolic processes, supporting growth and maintaining sucrose homeostasis. Up right, in response to stress, sucrose and T6P levels decrease, causing the dissociation of TPS-SnRK1 complexes and possibly the activation of other SnRK1 pools. This leads to high SnRK1 activity in the nucleus and strong activation of stress responses. Bottom left, the inability to perceive T6P disrupts sucrose homeostasis. In the absence of TPS5/6/7, a pool of non-nuclear SnRK1 is aberrantly active, promoting catabolic processes and growth repression despite high sucrose and T6P levels. The inhibition of other (non-nuclear) pools of SnRK1 by T6P or other signals/factors is normal. Bottom right, under stress, nuclear SnRK1 activity is weaker in the *tps5/6/7* mutant leading to mildly compromised stress responses. This is possibly caused by the overaccumulation of T6P and other sugars when TPS5/6/7 are depleted. White filling: inactive proteins, red or blue filling: active proteins.

Sensing and managing sucrose availability through T6P is crucial for maintaining sugar homeostasis and for coordinating growth with carbon availability (Baena-González and Lunn 2020; Fichtner and Lunn 2021). A tight coupling between sucrose and T6P exists across all tested plant species and conditions (Lunn et al. 2006; Martínez-Barajas et al. 2011; Wingler et al. 2012; Carillo et al. 2013; Martins et al. 2013; Nunes et al. 2013a; Wahl et al. 2013; Henry et al. 2014; Sulpice et al. 2014; Yadav et al. 2014; Zhang et al. 2015; Fichtner et al. 2020). This coupling was also evident in this study, where the levels of sucrose and T6P in Col-0 seedlings grown under control light were similar to those grown under high light (Fig. 2B and Table 1). However, this feedback mechanism appeared disrupted in the *tps5/6/7* mutant, where high light drove a prominent rise in T6P but this failed to maintain sucrose homeostasis, resulting in higher sucrose levels than in the control light condition. Our data suggests that this defective T6P-sucrose coupling is due to compromised T6P perception. First, the inability to sense T6P would be expected to impair sucrose utilization, resulting in an accumulation of sucrose that, in turn, stimulates further T6P synthesis. This would create a vicious cycle of sucrose/T6P production that worsens under conditions of increased carbon availability (e.g., under higher light intensities that promote photosynthesis or when exogenous sucrose is supplied). This is reminiscent of insulin resistance in animals, where tissues fail to respond to insulin, resulting in elevated blood sugar due to impaired glucose uptake and utilization. In response, the pancreas increases insulin production as a compensatory mechanism (James et al. 2021). The idea that *tps5/6/7* is compromised in T6P-perception is consistent with our T6P and sucrose measurements, with the root growth defects of the *tps5/6/7* mutant and with the inability of sugars to rescue these growth defects. Notably, the increase in sucrose under higher light intensities was accompanied by an even larger relative increase in T6P. This may indicate that not only sucrose and central metabolism are perturbed, but that the unknown regulatory loops that lead to an increase in T6P levels when sucrose rises are amplified in *tps5/6/7* seedlings. Second, given the reported increase in TCA cycle intermediates and amino acids caused by rising T6P in leaves (Figueroa et al. 2016; Ishihara et al. 2022; Avidan et al. 2024), the inability to sense T6P would be expected to have a negative impact on the levels of these central building blocks. This aligns with the broad depletion of amino acids, particularly BCAAs and Lys, observed in the *tps5/6/7* mutant (Fig. 2C and Supplementary Fig. 7).

What, then, causes this amino acid depletion? Possible explanations include (1) impaired nitrogen assimilation or *de novo* amino acid biosynthesis, (2) increased amino acid incorporation into proteins, (3) decreased protein turnover, and (4) enhanced amino acid catabolism. The first possibility seems less plausible, as the levels of Glu and Gln were either comparable between genotypes or slightly elevated in the *tps5/6/7* mutant (Supplementary Fig. 7). Additionally, the mutant accumulated certain amino acids (e.g., Pro and Ala) to higher levels (Supplementary Fig. 7), indicating that nitrogen assimilation and amino acid biosynthetic pathways remain functional. Alternatively, amino acid depletion could arise from altered dynamics in protein synthesis and degradation. However, total protein content did not differ significantly between genotypes (Supplementary Fig. 8). The most compelling explanation is that T6P insensitivity in the mutant reconfigures metabolism to promote the use of BCAAs and Lys as alternative substrates for ATP production. This is supported by several lines of evidence. First, the ratio of ATP synthesis to oxygen consumption is lower when amino acids are respired instead of glucose (Leverve et al. 2007). Second, the oxygen consumption rates of the mutant were >1.5-fold higher than in Col-0 both in the absence and presence of exogenous sucrose (3-6h interval). Furthermore, the positive effect of sucrose on respiration rates was significantly lower in the mutant than in Col-0 (6-24h interval). Third, the difference in amino acid levels between Col-0 and the *tps5/6/7* mutant was most prominent under high light conditions (Fig. 2C; Supplementary Fig. 7B), where Col-0 exhibited increased growth (Fig. 2A).

Intriguingly, in *tps5/6/7* seedlings, root growth was more severely impaired than that of shoots (Fig. 1C and Supplementary Fig. 4), although both mutant tissues were largely depleted of BCAAs (Fig. 2C and Supplementary Fig. 7B). This disparity may be due to the greater metabolic flexibility of green tissues, where photosynthetic activity can generate ATP as well as respiratory substrates such as TCA cycle intermediates. Roots, on the other hand, rely entirely on imported sucrose and on respiration for energy and growth (Hennion et al., 2019), rendering them more vulnerable to disruptions in sucrose signalling. This may explain why, in high light conditions, the levels of malate and aconitate (and to a lesser extent other organic acids) were higher in the shoots of the mutant compared to Col-0 whilst the opposite was true in root tips (Supplementary Fig. 7A).

Why is T6P perception compromised in the *tps5/6/7* mutant? The active site residues critical for T6P binding appear conserved in the TPP-like domain of TPS-II proteins (Lunn 2007). Furthermore, their interaction with SnRK1α1 is enhanced in a T6P-dependent manner (Fig. 4). We therefore postulate that TPS-II proteins act as T6P sensors and that, upon T6P binding, their interaction with SnRK1 is strongly enhanced. However, it is also possible that T6P binds to other proteins, including SnRK1 (Fig. 5). Indeed, T6P has been shown to bind to the SnRK1α1 subunit, preventing its interaction with the SnRK1 activating kinase (SnAK) and, thereby, SnRK1 T-loop phosphorylation and activity (Zhai et al. 2018; Blanford et al. 2024).

Regardless of which protein binds T6P, our results demonstrate that TPS-II proteins are crucial for proper SnRK1 regulation. In the absence of TPS5/6/7, high T6P fails to inhibit SnRK1, with this unchecked SnRK1 activity then causing the growth defects of the *tps5/6/7* mutant. Notably, a recent study showed that loss-of-function alleles of the SnRK1α and SnRK1 βγ subunits rescue both the flowering and embryogenesis defects of the *tps1-2* mutant (Zacharaki et al. 2022). This further supports the idea that a lack of T6P – real, as in *tps1-2*, or perceived, as in *tps5/6/7* seedlings – leads to aberrant SnRK1 activity that ultimately suppresses growth and development.

Our results also show that lack of TPS5/6/7 affects nuclear and non-nuclear SnRK1 pools differently (Fig. 5). On one hand, several lines of evidence suggest that, under favourable conditions, TPS-II proteins promote growth through inhibition of a distinct SnRK1 pool (Fig. 5); consequently, in the absence of TPS-II proteins, this SnRK1 pool becomes overactive, causing aberrant repression of growth. First, the marked growth defects of *tps5/6/7* are rescued by SnRK1 depletion (Fig. 3A). Second, TPS-II proteins interact physically with SnRK1 [Fig. 4; (Van Leene et al. 2022)] and this interaction is strongly stimulated by T6P (Fig. 4). Third, the T6P:suc ratios of the *tps5/6/7* mutant shoots (0.169, 1.6-fold higher than in Col-0, Table 1) are remarkably similar to those reported for rosettes of SnRK1α1 overexpressors at the end of the night [0.170, 1.4-fold higher than in Col-0, (Peixoto et al. 2021)], and clearly different from those of rosettes overexpressing an *E. coli* TPS (otsA) [2.831, 21-fold higher than in Col-0, (Yadav et al. 2014)]. Given that basal nuclear activity is not higher in the mutant (Fig. 3C), it is likely that the subcellular SnRK1 pool directly regulated by TPS-II proteins is non-nuclear. On the other hand, the stimulation of SnRK1 activity by energy stress appears weaker in the *tps5/6/7* mutant (Fig. 3C) suggesting that, without TPS-II proteins, SnRK1 is partly insensitive to energy signals. We envision two possible scenarios to explain this. Lower activation of SnRK1 by energy stress in the *tps5/6/7* mutant could be a direct consequence of TPS-II depletion. TPS-II proteins may sequester a non-nuclear pool of SnRK1 in repressor complexes that respond to energy signals (Fig. 5). In the absence of TPS5/6/7, this SnRK1 pool would be aberrantly active, yet partly insensitive to further activation by energy stress, as it lacks the proteins that associate SnRK1 to the energy signal. This would be reminiscent of how SnRK2 proteins regulate SnRK1 in response to abscisic acid (ABA) (Belda-Palazón et al. 2020, 2022). However, contrary to SnRK2s (Belda-Palazón et al. 2020, 2022), we could not detect TPS7-mCherry in the nucleus at EN (Supplementary Fig. 10). This suggests that TPS-II proteins form non-nuclear SnRK1 repressor complexes, which is consistent with the fact that transient co-expression of TPS-II proteins with SnRK1 prevents its nuclear localisation and activity (Van Leene et al. 2022). It is therefore difficult to explain how the lack of TPS5/6/7 could by itself reduce the presence of SnRK1 in the nucleus. A second scenario is that defective activation by energy stress might be an indirect consequence of TPS-II depletion. Aberrantly high T6P levels could interfere with T-loop phosphorylation by the SnAK kinase (Zhai et al. 2018; Blanford et al. 2024). However, the levels of T-loop phosphorylation were not significantly different between Col-0 and *tps5/6/7* seedlings (Supplementary Fig. 9), arguing against this possibility. Alternatively, the impact on SnRK1 localization and activation by energy stress could be an indirect consequence of the accumulation of other sugars in the mutant through T6P/TPS-independent mechanisms yet to be resolved. Distinguishing between a direct and an indirect effect of TPS-II proteins on SnRK1 activation by energy stress would require uncoupling elevated T6P and soluble sugars from TPS5/6/7 depletion in the future.

Despite these disruptions, the *tps5/6/7* mutant retains some capacity for growth, suggesting that additional factors contribute to SnRK1 regulation. Indeed, our data show that SnRK1 interacts with most TPS-II proteins (Fig. 4C), supporting previous findings that the entire TPS-II family participates in SnRK1 regulation (Van Leene et al. 2022). Moreover, TPS-IIs exhibit both distinct and overlapping expression patterns (Ramon et al. 2009; Mergner et al. 2020), suggesting that TPS-II proteins may play partly redundant functions and that TPS8-11 may to some extent compensate for the loss of TPS5-7. Nevertheless, the extent to which other TPS-II proteins contribute to SnRK1 regulation and growth requires further investigation.

In summary, our findings uncover a crucial role for TPS5/6/7 as mediators of the sucrose status, fine-tuning SnRK1 activity to balance metabolism and growth. Further studies are needed to fully elucidate the roles of TPS-II proteins across different tissues and environmental conditions, as well as their interactions with other components of the T6P-SnRK1 regulatory network. Given their conservation across plant species, such insights will not only deepen our understanding of plant energy sensing but also inform strategies to optimize crop performance under increasingly challenging environments.

## SUPPLEMENTARY FIGURE LEGENDS

**Supplementary Figure 1. Expression atlas of class II TPS proteins in Arabidopsis.** Relative protein abundance in different tissues based on data from Mergner et al. (2020) and visualized with ATHENA proteomics (https://athena.proteomics.wzw.tum.de/master_arabidopsisshiny/). *RKD2*, egg-cell like callus; *CIM*, callus; *CC3*, root cell culture early; *CC10*, root cell culture late; *SP*, sepal (flower stage 15); *PT*, petal (flower stage 15); *ST*, stamen (flower stage 15); *CP*, carpel (flower stage 15); *SQ*, silique (stage 3); *FL*, flower (stage 15); *SQSP*, silique septum; *SQV*, silique valves; *CLLF*, 1^st^ cauline leaf; *LFD*, rosette leaf 7, distal part; *LFP*, rosette leaf 7, proximal part; *LFPT*, rosette leaf 7, petiole; *SCLF*, senescent leaf; *CT*, cotyledons; *SAMCT*, shoot apical meristem, cotyledons and first leaves; *P*, pollen; *RT*, root; *RTP*, root tip; *RTUZ*, root upper zone; *EB*, seed (embryo stage 10); *SD*, seed (mature, dry); *SDIMB*, seed (mature, imbibed); *ND*, stem, 1^st^ node; *IND*, stem, 2^nd^ internode; *FLPD*, flower pedicle (flower stage 15); *HY*, hypocotyl.

**Supplementary Figure 2. Generation of *tps* mutants.** (**A**) Schematic representation of the insertion sites of the *tps5*, *tps6*, and *tps7* T-DNA mutants used in this study. The ATG start codon and TAA/TGA stop codons are indicated. Exons are shown as black boxes, introns as black lines, and T-DNA insertion sites as triangles above the corresponding gene diagrams. Scale bar: 400 bp. (**B**) Confirmation of *tps* mutant identities by genotyping using T-DNA left border and gene-specific primers (LP and RP, indicated in **A**). (**C**) RT-PCR analyses of the *TPS6* (upper panel) and *TPS7* (lower panel) transcripts in *tps6-1* and *tps7-1* mutants, respectively. Amplification of *ACTIN* mRNA was used as a control.

**Supplementary Figure 3. Phenotype of single *tps5*, *tps6*, and *tps7* mutants.** Left, representative images of 14d-old Col-0, *tps5*, *tps6* and *tps7* seedlings grown vertically on 0.5× MS medium for 7 d and transferred to 0.5× MS medium for 7 d. White marks indicate root length at transfer. Scale bar 1 cm. Right, quantification of primary root (PR) length, measured from the white mark to the root tip (*n*=3 independent experiments, 48-64 seedlings per genotype). In the violin plot, dotted lines represent the quartiles and the dashed line, the median. Different letters indicate statistically significant differences between genotypes (*p*<0.05, one-way ANOVA with Tukey HSD test).

**Supplementary Figure 4. Rosette phenotype of Col-0 and *tps5/6/7* plants.** Representative images of rosettes from Col-0 (left) and *tps5/6/7* (right) plants grown on soil under long-day conditions (16 h light, 110 μmol m^−2^ s^−1^, 22 °C / 8 h dark, 18 °C). Images were captured over 12 days, from day 17 to day 29 after sowing.

**Supplementary Figure 5. Impact of glucose and fructose on the *tps5/6/7* PR growth phenotype.** Left, representative images of 14d-old Col-0 and *tps5/6/7* seedlings grown vertically on 0.5× MS medium for 7 d and transferred to 0.5× MS medium supplemented (right) or not (left) with 30 mM Glucose + 30 mM fructose for 7 d. White marks indicate root length at transfer. Scale bar 1 cm. Right, quantification of PR length, measured from the white mark to the root tip (n*=*1 independent experiment, 22-24 seedlings per genotype per condition). In the violin plot, dotted lines represent the quartiles and the dashed line, the median. Different letters indicate statistically significant differences between genotype and condition combinations (*p*<0.05, one-way ANOVA with Tukey HSD test).

**Supplementary Figure 6. Impact of TPS5/6/7 depletion on sugars related to sucrose and trehalose metabolism.** Levels of sucrose 6-phosphate, glucose, fructose, and α,α trehalose in the indicated tissues of 14d-old Col-0 and *tps5/6/7* seedlings. Seedlings were grown vertically on 0.5× MS medium under control light conditions (110 μmol m^−2^ s^−1^) for 7 d, transferred to fresh 0.5× MS plates and grown for 7 d under control light (CL; left, white) or high light conditions (HL; 200 μmol m^−2^ s^−1^; right, yellow). Graphs show the mean of 3 independent experiments (each consisting of 200-250 seedlings per genotype per condition, error bars, SD). Asterisks indicate statistically significant differences between genotypes within each tissue and condition (*p*<0.05, two-tailed unpaired Student t-test).

**Supplementary Figure 7. Impact of TPS5/6/7 depletion on primary metabolism.** (**A**) Ratios of primary metabolites in the indicated tissues of 14d-old Col-0 and *tps5/6/7* seedlings. Seedlings were grown vertically on 0.5× MS medium under control light conditions (110 μmol m^−2^ s^−1^) for 7 d, transferred to fresh 0.5× MS plates and grown for 7 d under control light (CL; left) or high light conditions (HL; 200 μmol m⁻² s⁻¹; right). Colors indicate the log2 ratios of the mean values (*tps5/6/7 vs*. Col-0; *n*=3). Asterisks indicate statistically significant changes in metabolite levels between genotypes within each tissue and light condition (*\*p*<0.05, *\*\*p*<0.01, *\*\*\*p*<0.001; two-tailed unpaired Student and Welch’s t-tests). (**B**) Levels of Lysine and Valine in the indicated tissues of 14d-old Col-0 and *tps5/6/7* seedlings grown as in (**A**) under contrasting light intensities (CL, white; HL, yellow). Graphs show the mean of 3 independent experiments (each consisting of 200-250 seedlings per genotype per condition, error bars, SD). Asterisks indicate statistically significant differences between genotypes within each tissue and condition (*p*<0.05, two-tailed unpaired Student and Welch’s t-tests).

**Supplementary Figure 8. Impact of TPS5/6/7 depletion on total protein content.** Total protein levels in the indicated tissues of 14d-old Col-0 and *tps5/6/7* seedlings. Seedlings were grown vertically on 0.5× MS medium under control light conditions (110 μmol m^−2^ s^−1^) for 7 d, transferred to fresh 0.5× MS plates and grown for an additional 7 d under control light (CL; left) or high light conditions (HL; 200 μmol m^−2^ s^−1^; right). Samples were harvested at the end of the night. Graphs show the mean of 3 independent experiments (each consisting of 200-250 seedlings per genotype per condition, error bars, SD). Asterisks indicate statistically significant differences between genotypes within each tissue and condition (*p*<0.05, two-tailed unpaired t-test).

**Supplementary Figure 9. Impact of TPS5/6/7 depletion on SnRK1ɑ1 T-loop phosphorylation.** (**A**) Representative immunoblots showing SnRK1α T-loop phosphorylation (pSnRK1α1 and pSnRK1α2) and total SnRK1α1 levels detected, respectively, with anti-phosphoAMPK (p-AMPK) and anti-SnRK1α1 antibodies. Proteins were extracted from 14d-old Col-0 and *tps5/6/7* seedlings, either maintained in the light (L) or transferred to darkness (D) at ZT2 for 4 h and harvested at ZT6. (**B**) Quantification of SnRK1ɑ1 T-loop phosphorylation, presented as the pSnRK1α1/SnRK1α1 ratio. Bars show the mean of 4 independent experiments (error bars, SD). Asterisks indicate significant differences between genotypes (*p*<0.05; two-tailed unpaired Student t-test).

**Supplementary Figure 10. Subcellular localization of TPS7-mCherry.** Representative fluorescence microscopy images of the root meristem of Arabidopsis seedlings stably expressing *TPS7-mCherry* (line C#2). Scale bar: 25 µm.

**Supplementary Figure 11. Experimental set-up for assessing the TPS7-SnRK1ɑ1 interaction.** Cleared protein extracts from 14d-old *proTPS7::TPS7-mCherry* (**A**) or *proSnRK1α1::SnRK1α1-GFP* (**B**) seedlings, harvested at EN, were aliquoted and incubated with ChromoTek RFP- or GFP-Trap® Magnetic Agarose beads, respectively, in the absence (H₂O control) or presence of the indicated concentrations of T6P. (**A**) Immunoprecipitated mCherry-tagged TPS7 and co-immunoprecipitated SnRK1α1 were assessed by immunodetection (Fig. 4A-B). (**B**) Proteins interacting with GFP-tagged SnRK1α1 were identified by MS/MS analyses (Fig. 4C).

## ACKNOWLEDGEMENTS

We thank Umesh Yadav and Andrzej Koczut for the identification of the single *tps5*, *tps6*, and *tps7* mutants and the generation of the *tps5/6/7* mutant, respectively. Vera Nunes is thanked for excellent plant management and Jenny Bormann, Svenja Heimann and Andrea Leisse for excellent technical support with LC-MS/MS (JB, SH) and GC/MS analyses (AL). This work was supported by Fundação para a Ciência e a Tecnologia grants UIDB/04551/2020 (DOI 10.54499/UIDB/04551/2020), UIDP/04551/2020 (DOI 10.54499/UIDP/04551/2020, GREEN-IT-Bioresources for Sustainability), LA/P/0087/2020 (LS4FUTURE Associated Laboratory, DOI: 10.54499/LA/P/0087/2020), PTDC/BIA-FBT/4942/2020 (DOI: <u>10.54499/PTDC/BIA-FBT/4942/2020</u>), individual work contract 2023.06360.CEECIND/CP2836/CT0009 (AC, DOI: 10.54499/2023.06360.CEECIND/CP2836/CT0009), and PhD fellowship (DR-B, PD/BD/150239/2019) and by the Max Planck Society (CC, JEL and MS). For the work carried in Oxford, we thank the John Fell Fund (0013575) for funding critical equipment.

## MATERIALS AND METHODS

A list of all plant lines, primers, and antibodies used in this study is provided in Supplementary Tables S2a-S2c.

### Plant material and growth conditions

All *Arabidopsis thaliana* plants used in this study are in the Columbia (Col-0) background. The *tps5* T-DNA insertion mutant (SALK_144791, *tps5-1*) was previously described (Tian et al., 2019). The *tps6* (GK-362A03) and *tps7* (GK_341E02) mutants were identified from the publicly available collection of *Arabidopsis thaliana* T-DNA insertion lines (http://signal.salk.edu/cgi-bin/tdnaexpress). Homozygosity of the *tps* mutations was confirmed by genomic PCR (Supplementary Fig. 2), using LP, RP and T-DNA LB primers (LBb1.3 and o8409 for SALK and GABI-Kat lines, respectively; Supplementary Table S2a) (Koczut et al., manuscript in preparation). The loss of full-length transcripts was verified by RT-PCR (Supplementary Fig. 2). The *snrk1α1-3* (Mair et al. 2015), *sesquiα2-2* [*snrk1α1-3*^-/-^*snrk1α2-2^+/-^*; referred as *sesqui*; (Belda-Palazón et al. 2020)], *pro35S:NLS-ratACC-GFP-2xHA* (referred as *NLS-ACC*; (Muralidhara et al. 2021)) and *proSnRK1α1::SnRK1α1-GFP* (Crozet et al., 2016) have been previously described. The *tps5/7* and *tps5/6/7* mutants were obtained by crossing *tps5-1* with *tps7* (GK_341E02) and subsequently with *tps6* (GK-362A03) for the triple mutant. F1 seeds were grown and allowed to self-fertilize to produce the F2 generation. Homozygosity of individual plants from a segregating F2 population was confirmed by PCR using gene-specific and T-DNA primers (Supplementary Table S2b; Koczut et al., manuscript in preparation). Combinatorial *tps5/6/7/snrk1α1-3*, *tps5/6/7/sesqui*, *tps5/6/7/ NLS-ACC* and *tps5/6/7/proSnRK1α1::SnRK1α1-GFP* mutants were obtained by crossing the *tps5/6/7* mutant with *snrk1α1-3*, *sesquiα2-2*, *pro35S:NLS-ratACC-GFP-2xHA* or *proSnRK1α1::SnRK1α1-GFP*, respectively.

Seeds were surface sterilized for 1 min in 70% (V/V) ethanol followed by 10 min in 20% (V/V) bleach under gentle rocking and then washed at least 5 times with sterile water. Sterilized seeds were stratified in the dark at 4°C for 2-3 days. Unless otherwise specified, plants were grown under long-day conditions (16 h light, 110 μmol m^−2^ s^−1^, 22 °C / 8 h dark, 18 °C) on 0.5× MS medium (0.05% MES and 0.8% phytoagar).

### Molecular cloning and Arabidopsis transformation

For generating the *pTPS7::TPS7-mCherry* complementation lines, relevant DNA fragments were amplified by PCR using Phusion™ High-Fidelity DNA Polymerase and primers containing appropriate overhangs (Supplementary Table S2b). Fragments were thereafter assembled into the final vector using the Gibson Assembly® method. The full-length *TPS7* gene (AT1G06410, 2738 bp), a 2000 bp region upstream of the *TPS7* start codon (promoter), and a downstream terminator sequence (833 bp) were separately amplified from Arabidopsis Col-0 genomic DNA. The mCherry sequence (708 bp) was amplified from a construct for expressing a Golgi-mCherry marker (Nelson, 2007). A 5039 bp segment of the pCB302 binary vector (Xiang et al. 1999)sequence, excluding the C-terminal GFP tag region, was amplified for the construct backbone. Purified PCR products were incubated at 50°C for 1h with the NEBuilder® Hifi DNA assembly master mix, following the manufacturer’s instructions. The resulting constructs were introduced into *E. coli* (TOP10) cells, and correct assembly was validated by colony PCR and sequencing. Verified constructs were introduced into *Agrobacterium tumefaciens* GV3101, and *tps7* (GK_341E02) plants were transformed by the floral dip method (Clough & Bent, 1998). BASTA®-resistant transformants were selected based on their segregation ratio (T2) and homozygosity (T3).

### Phenotyping assays

For root growth assays, seedlings were grown vertically for 7 days on 0.5× MS medium under control light conditions (16 h light, 110 μmol m^−2^ s^−1^, 22°C / 8 h dark, 18°C). Seedlings with comparable root sizes were then transferred to fresh 0.5× MS plates, supplemented with sugars when indicated (30 mM sucrose or 30 mM fructose and 30 mM glucose). When indicated, fresh 0.5× MS plates were shifted to high light (16 h light, 200 μmol m^−2^ s^−1^, 22°C / 8 h dark, 18°C). The position of the root tip was marked immediately after transfer and seedlings were grown on the fresh plates for an additional 7 days before scanning. Post-transfer root growth was measured as primary root (PR) length from scanned plates. Tracings and measurements were performed using the NeuronJ plugin (Meijering et al. 2004) in ImageJ. In assays involving segregating *sesqui* and *tps5/6/7/sesqui* lines, seedlings were harvested after scanning the plates, and their genotype was determined by PCR.

### Metabolite extraction and measurement

For metabolite analyses, Col-0 and *tps5/6/7* seedlings were grown vertically for 7 days on 0.5× MS medium under control light conditions. Seedlings with comparable root sizes were then transferred to fresh 0.5× MS plates, where they grew on calibrated Nitex mesh (pore size 30 µm) under control (16 h light, 110 μmol m^−2^ s^−1^, 22°C / 8 h dark, 18°C) or high light conditions (16 h light, 200 μmol m^−2^ s^−1^, 22°C / 8 h dark, 18°C) for an additional 7 days. 14d-old shoots, roots (excluding the apical region) and root apices (∼0.5 cm) were harvested separately at the end of the night (EN), flash frozen and reduced to a fine powder in liquid nitrogen. For LC-MS/MS- and GC-MS-based quantifications, approximately 18 mg of finely ground plant material was quenched in a 3:7 (v/v) chloroform:methanol mixture at liquid nitrogen temperature and warmed to −20 °C for 2 h. The chloroform organic phase was subsequently extracted twice with ice-cold ultra-pure water. Both water phases were combined and dried to a powder in a centrifugal vacuum dryer at 20 °C. Pellets were resuspended in 350 µL of ultra-pure water and filtered through a Multiscreen Ultracel-10 (Millipore) membrane to eliminate high molecular weight components. Dilutions of 1:10 were used to quantify T6P, other phosphorylated intermediates and organic acids via high-performance anion-exchange chromatography coupled to tandem mass spectrometry, as described in (Lunn et al. 2006) with modifications (Figueroa et al. 2016). Soluble sugars and sugar alcohols were quantified by high-performance hydrophilic interaction liquid chromatography (HILIC) on a 150 x 2.1 mm Luna Omega Sugar column (3 µm, 100Å; Phenomenex Inc.; Aschaffenburg, Germany; www.phenomenex.com) column fitted with a SUGAR 4 x 2.0 mm Security Guard Cartridge (Phenomenex), coupled to a QTrap 5500 triple quadrupole mass spectrometer (AB Sciex, Foster City, USA; sciex.com). The column was maintained at 25 °C and equilibrated with eluent A (90% acetonitrile, 5% isopropanol, 5% water, by volume) before injection of samples (1 μL). Samples were spiked with ^13^C-labelled internal standards for correction of ion suppression and other matrix effects. Sugars were eluted (flow rate 0.25 mL min^-1^) by mixing eluent A and eluent B (5% acetonitrile, 25% isopropanol, 70% water, by volume) with the following gradient:: 0 – 2.5 min [10 % B]; 2.5 – 25 min [10 – 30 % B]; 25 – 30.5 min [30 % B]; 30.5 – 31 min [30 – 10 % B]; 31 – 40 min [10 % B]. Sugars were quantified by tandem mass spectrometry in multiple reaction mode (MRM) using the settings and stable-isotope-labelled internal standards (SIL-IS) summarized in Supplementary Tables S3a and S3b.

Total protein amounts were extracted from the left-over pellets of the chloroform:methanol extraction (Hendriks et al. 2003). Pellets were dried at room temperature, resuspended in 400 µL NaOH, shaken vigorously, heated at 120 °C for 30 min, cooled down at room temperature and centrifuged at 21,000 *g* for 10 min at room temperature. Total protein amounts were determined with the Bradford method, using bovine serum albumin as standard.

Amino acids and other polar metabolites were measured by drying 200 µL of the organic phase and derivatizing it according to (Roessner et al. 2001). 1 µL of the derivatized sample was analyzed on a Combi-PAL autosampler (Agilent Technologies) coupled with an Agilent 7890 GC and Leco Pegasus 2 TOF-MS (LECO, St. Joseph, MI, USA) (Lisec et al. 2006). Chromatograms were exported to R software via Leco CHROMA TOF (v. 3.25) for peak detection, retention time alignment, and library matching using R/TARGETSEARCH (Cuadros-Inostroza et al. 2009). Metabolites were quantified based on the peak intensity of a selective mass and normalized to FW and total ion count. Final values were normalized by the median intensity of the metabolite across all samples.

### Measurements of oxygen consumption rates

Oxygen consumption in Arabidopsis seedlings was measured using Screw Cap Septum Vials (37.5 mL). Ten-day-old seedlings, grown on vertical 0.5× MS plates, were transferred into vials partially filled with 20 mL of 0.5× MS medium, with or without 1% sucrose, and solidified with 0.8% agar. Each vial contained approximately 15–20 seedlings. Incubation and measurements were carried out at 22 °C in darkness (under green light) to prevent photosynthesis and the resulting oxygen production. Seedlings were transferred at ZT4. Once the transfer was completed, the vials were quickly closed and T0 oxygen measurements were initiated. Each measurement lasted 25 seconds per sample. Oxygen concentration in the air space was monitored at three time points (3, 6 and 24 hours) using a needle-type oxygen sensor (Unisense), which was inserted through the septum to measure the oxygen concentration. The decrease in oxygen levels between 3 and 6h and between 6 and 24h was used as a proxy for respiration rate in the corresponding time interval. Following the measurements, the fresh weight of the seedlings and the headspace volume of each vial were recorded for normalization to express respiration rates as µmol O_2_ consumed /g FW /hour.

### Subcellular localization analyses by confocal microscopy

SnRK1α1 localization was observed at the end of the night in roots of SnRK1α1-GFP and *tps5/6/7*;SnRK1α1-GFP seedlings grown vertically on 0.5× MS plates under long-day conditions (16 h light, 110 μmol m^-2^ s^-1^, 22°C / 8 h dark, 18°C) for 8 days. On day 8, one hour before dawn, roots were stained for 2 min with an aqueous solution of propidium iodide (PI; 10 μg/mL) and images were acquired on a Zeiss AxioObserver 780, using a 40× water immersion objective. For the visualization of GFP, pinholes were adjusted to 1 Airy Unit (488 nm/500-530 nm). For quantitative analysis of GFP, the 488 nm laser power was set to 3.0% transmission, and the master gain was set to 800. For the visualization of cell walls, pinholes were also adjusted to 1 Airy Unit (561 nm/600-660 nm). Post-acquisition image processing was performed using ImageJ (http://rsb.info.gov/ij/).

The nuclear/cytoplasmic localization of SnRK1α1 was assessed using CLSM and ImageJ software, calculating the N/C ratio (N/C = Mean Nuclear Fluorescence Intensity / Mean Cytoplasmic Fluorescence Intensity, with Mean Fluorescence Intensity being the ratio between the total fluorescence intensity measured and the number of pixels measured) (Kelley and Paschal, 2019). The area of the nucleus was determined by analyzing the bright field acquired through the transmitted light mode whilst the area of the cytoplasm was selected using the PI signal as a reference.

TPS7 localization was observed at the end of the night in roots of *TPS7-mCherry* seedlings (line C#2), grown vertically on 0.5× MS plates under long-day conditions (16 h light, 110 μmol m^-2^ s^-1^, 22°C / 8 h dark, 18°C) for 8 days. On day 8, one hour before dawn, roots were mounted in water. Images were acquired using a Leica Stellaris 8 system, using a 63x water immersion objective. For the visualization of mCherry, a white light laser was used at 85% power and 8% intensity to excite the fluorophore at 581 nm. Red fluorescent signal was detected on the 587 – 700 nm range, with a hybrid HyD S detector operating in analogue mode at 600 Hz. Pinholes were kept at 1 airy units (AU), and the final image was acquired by averaging 6x over each line and accumulating signal 2x over the entire frame. Gray value histograms were adjusted, and scale bars were added to the images in Leica’s software Las X Office v. 1.4.7.28982, prior to being exported.

### In planta SnRK1 activity assay

Arabidopsis *NLS-ACC* and *tps5/6/7/NLS-ACC* seedlings were grown vertically for 7 days on 0.5× MS medium. Seedlings with comparable root sizes were then transferred to fresh 0.5× MS plates and grown for an additional 7 days. On day 14, seedlings were either kept in the light or shifted to darkness at ZT2 for 4 hours. Seedlings of both groups were harvested at ZT6, flash-frozen and reduced to a fine powder in liquid nitrogen. Proteins were extracted on ice with extraction buffer [1.5 mL buffer/ g fresh weight; 50 mM Tris-HCl pH 8.0, 150 mM NaCl, 1 mM EDTA, 1% (v/v) Triton X-100, 3 mM DTT, 50 μM MG-132, 0.002% (v/v) phosphatase inhibitor cocktail #2 (Sigma P572G), 0.002% (v/v) phosphatase inhibitor cocktail #3 (Sigma P0044) and cOmplete™ protease inhibitor cocktail (Roche; 1 tablet/10mL)]. Homogenates were cleared by centrifugation at 21130 *g* for 15 min at 4 °C, and protein concentration in the resulting supernatants was quantified using the Bradford assay. Proteins were denatured in Laemmli buffer, loaded onto 10% acrylamide gels and resolved by SDS-PAGE. Separated proteins were transferred to PVDF membranes using transfer buffer (192 mM glycine, 25 mM Tris, 0.1% SDS, 20% ethanol) and a Bio-Rad wet blotting transfer system (90 min at 4°C, 80 V). Membranes were blocked for 1 hour (5% w/v non-fat dry milk in 1X TBS, 0.05% Tween®) and probed with anti-P(S79)-ACC or anti-GFP (Supplementary Table S2c) under gentle rocking overnight at 4 °C. Anti-rabbit IgG or anti-mouse IgG were used as secondary antibodies. Protein band intensities were determined in ImageJ (version 2.1.0), and p-ACC/GFP ratios were analyzed with GraphPad Prism (version 10.3.1).

### SnRK1 T-loop phosphorylation

Arabidopsis Col-0 and *tps5/6/7* seedlings were grown and treated as described in the section “In planta SnRK1 activity”. Protein extraction, SDS-PAGE and immunoblotting were likewise performed as described in that section. Membranes were probed with anti-P(T172)-AMPKα or anti-SnRK1α1 (Supplementary Table S2c). Anti-rabbit IgG was used as secondary antibody. Protein band intensities were determined in ImageJ (version 2.1.0), and p-SnRK1α1/SnRK1α1 ratios were analyzed with GraphPad Prism (version 10.3.1).

### Coimmunoprecipitation experiments

For assessing the interaction of TPS7 with SnRK1α1, 14d-old *proTPS7::TPS7-mCherry* seedlings (line C #2) grown vertically on 0.5× MS medium were harvested at the end of the night, flash frozen and reduced to a fine powder in liquid nitrogen. Proteins were extracted on ice with immunoprecipitation (IP) buffer [1.5 mL buffer/ g fresh weight; 50 mM Tris-HCl pH 8.0, 150 mM NaCl, 1 mM EDTA, 1% (v/v) Triton X-100, 3 mM DTT, 50 μM MG-132, 0.002% (v/v) phosphatase inhibitor cocktail #2 (Sigma P572G), 0.002% (v/v) phosphatase inhibitor cocktail #3 (Sigma P0044) and cOmplete™ protease inhibitor cocktail (Roche; 1 tablet/10mL)]. The resulting cleared protein extract was divided into 250 µL aliquots, to which 10 µl of one of the following was added to achieve the indicated concentrations in a final volume of 500µl: H_2_O, T6P (10 µM, 100 µM or 1 mM), UDPG (1 mM or 10 mM), trehalose (1 mM or 10 mM), G1P (10 mM), G6P (10 mM) or sucrose (10 mM). Volumes were then adjusted to 500 µL by adding 240 µL of dilution buffer [50 mM Tris-HCl pH 8.0, 150 mM NaCl, 1 mM EDTA, 3 mM DTT, 0.002% (v/v) phosphatase inhibitor cocktail #2 (Sigma P572G), 0.002% (v/v) phosphatase inhibitor cocktail #3 (Sigma P0044) and cOmplete™ protease inhibitor cocktail (Roche; 1 tablet/10mL)]. mCherry-tagged TPS7 was immunoprecipitated using ChromoTek RFP-Trap® Magnetic Agarose beads. Immunoprecipitated TPS7-mCherry was eluted in 50 µL of 2x Laemmli buffer by boiling for 5 min at 95 °C, and coimmunoprecipitated proteins were analyzed by western blotting using anti-RFP and anti-SnRK1α1 antibodies. Protein band intensities were determined in ImageJ (version 2.1.0), and SnRK1α1/TPS7-mCherry ratios were analyzed with GraphPad Prism (version 10.3.1).

To explore the effect of T6P on the SnRK1α1 interactome, 14d-old *proSnRK1α1::SnRK1α1-GFP* seedlings grown vertically on 0.5× MS medium were harvested at the end of the night, flash frozen and reduced to a fine powder in liquid nitrogen. Cleared protein extracts were prepared as described above, split into 300 µL aliquots, and mixed with either H_2_O or 1 mM T6P along 190 µL of dilution buffer to reach a final reaction volume of 500 µL. GFP-tagged SnRK1α1 was immunoprecipitated using ChromoTek GFP-Trap® Magnetic Agarose beads.

### Preparation of protein samples for LC-MS/MS

Samples for proteomic analysis were prepared using the single-pot, solid-phase-enhanced sample-preparation (SP3) strategy (Hughes et al. 2019). All buffers and solutions were prepared with mass spectrometry (MS)-grade water (Avantor, Radnor, PA, USA). Proteins from SnRK1α1-GFP pull down experiments were eluted twice in 50 µL of 2× elution buffer [2% (w/v) SDS, 20 mM TCEP, 80 mM chloracetamide, 100 mM HEPES pH 8.0] by heating each time for 5 min at 90 °C and clearing by centrifugation. The eluates were combined (100 µl in total) in a fresh tube, flash-frozen in liquid nitrogen, and stored at −80 °C. For LC-MS/MS analyses, eluted proteins were transferred to a 96-well plate. Then, 3 µl of a 50 µg µl^-1^ 1:1 mixture of hydrophilic (#45152105050250) and hydrophobic (#65152105050250) carboxylate modified Sera-Mag™ SpeedBeads (Cytiva, Marlborough, MA, USA) that were washed twice with MS-grade water were added to the samples. Afterwards, the samples were mixed shortly (1 min, 1000 rpm, RT) and collected by short centrifugation (10 sec, 200 ×g, RT). Protein binding was induced by the addition of an equal volume of pure ethanol (24 °C, 10 min, 1000 rpm), the beads were collected by a brief centrifugation step (10 sec, 200 ×*g*, RT) and the plate was placed on a magnetic stand. Beads were allowed to bind for at least 5 min before the supernatant was removed. The beads were then taken up in 180 µL 80% ethanol and transferred to a fresh multiwell plate. Subsequently the beads were washed four times with 180 µl 80% (vol/vol) ethanol prior to the addition of 100 µL digestion enzyme mix (0.6 μg of trypsin (V5111; Promega, Madison, WI, USA) and 0.6 µg LysC (125-05061; FUJIFILM Wako Pure Chemical, Osaka, Japan) in 25 mM ammonium bicarbonate). Samples were incubated at 37 °C for 19 h while shaking (1300 rpm). Next day, the samples were briefly centrifuged (10 sec, 200 ×*g*, RT) and placed on a magnet for 5 min. The clear solution containing the tryptic peptides was transferred to a fresh multiwell plate. The beads were taken up in 47 µL 25 mM ABC and incubated while shaking (RT, 10 min, 1000 rpm). The plate was then placed once more on a magnetic stand and after 5 min the cleared supernatant was again collected and combined with the recovered first peptide mix. This was followed by the addition of formic acid (FA; 2% final concentration) to the samples (trypsin inactivation).

### Sample clean-up for LC-MS/MS

Peptides were desalted on home-made C18 StageTips (Rappsilber et al. 2007) containing two layers of an octadecyl silica membrane (CDS Analytical, Oxford, PA, USA). All centrifugation steps were carried out at room temperature. The StageTips were first activated and equilibrated by passing 50 μL of methanol (600 × g, 2 min), 80% (v/v) acetonitrile (ACN) with 0.5% (v/v) FA (600 × g, 2 min) and 0.5% (v/v) FA (600 × g, 2 min) over the tips. Next, the acidified tryptic digests were passed over the tips (800 × g, 3 min). The immobilized peptides were then washed with 50 μL and 25 μL 0.5% (v/v) FA (800 × g, 3 min). Bound peptides were eluted from the StageTips by application of two rounds of 25 μL 80% (v/v) ACN with 0.5% (v/v) FA (800 × g, 2 min). After elution from the StageTips, the peptide samples were dried using a vacuum concentrator (Eppendorf, Hamburg, Germany) and the peptides were dissolved in 15 μL 0.1% (v/v) FA prior to analysis by MS.

### LC-MS/MS Analysis

LC-MS/MS analysis of peptide samples were performed on an Orbitrap Fusion Lumos mass spectrometer (Thermo Scientific, Waltham, MA, USA) coupled to a Easy nLC 1200 ultra high-performance liquid chromatography (UHPLC) system (Thermo Scientific, Waltham, MA, USA) that were operated in the one-column mode. The analytical column was a fused silica capillary (inner diameter 75 μm, outer diameter 360 µm, length 28 cm; CoAnn Technologies, Richland, WA, USA) with an integrated sintered frit packed in-house with Kinetex 1.7 μm XB-C18 core shell material (Phenomenex, Aschaffenburg, Germany). The analytical column was encased by a PRSO-V2 column oven (Sonation, Biberach, Germany) and attached to a nanospray flex ion source (Thermo Scientific, Waltham, MA, USA). The column oven temperature was set to 50 °C during sample loading and data acquisition. The LC was equipped with two mobile phases: solvent A (2% ACN and 0.2% FA, in water) and solvent B (80% ACN and 0.2% FA, in water). All solvents were of UHPLC grade (Honeywell, Charlotte, NC, USA). Peptides were directly loaded onto the analytical column with a maximum flow rate that would not exceed the set pressure limit of 950 bar (usually around 0.5 – 0.6 μL min^−1^) and separated on the analytical column by running a 105 min gradient of solvent A and solvent B at a flow rate of 300 nL/min (start with 3% (v/v) B, gradient 3% to 6% (v/v) B for 5 min, gradient 6% to 29% (v/v) B for 70 min, gradient 29% to 42% (v/v) B for 15 min, gradient 42% to 100% (v/v) B for 5 min and 100% (v/v) B for 10 min).

The mass spectrometer was controlled by the Orbitrap Fusion Lumos Tune Application and operated using the Xcalibur software. The MS settings for the different experiments are provided in Supplementary Table S4.

### Data processing and analysis

RAW spectra were submitted to an Andromeda (Cox et al. 2011) search in MaxQuant (2.0.3.0) using the default settings (Cox and Mann 2008). Label-free quantification and match-between-runs was activated (Cox et al. 2014). The MS/MS spectra data were searched against the Uniprot *A. thaliana* reference proteome UP000006548_3702_20231009_OPPG.fasta (one protein per gene; 27449 entries) and the project specific database ACE_0796_SOI_v01.fasta (2 entries). All searches included a contaminants database search (as implemented in MaxQuant, 245 entries). The contaminants database contains known MS contaminants and was included to estimate the level of contamination. Andromeda searches allowed oxidation of methionine residues (16 Da) and acetylation of the protein N-terminus (42 Da). Carbamidomethylation on Cystein (57) was selected as static modification. Enzyme specificity was set to “Trypsin/P”. The instrument type in Andromeda searches was set to Orbitrap and the precursor mass tolerance was set to ±20 ppm (first search) and ±4.5 ppm (main search). The MS/MS match tolerance was set to ±0.5 Da. The peptide spectrum match FDR and the protein FDR were set to 0.01 (based on target-decoy approach). For protein quantification unique and razor peptides were allowed. Modified peptides were allowed for quantification. The minimum score for modified peptides was 40. Label-free protein quantification was switched on, and unique and razor peptides were considered for quantification with a minimum ratio count of 2. Retention times were recalibrated based on the built-in nonlinear time-rescaling algorithm. MS/MS identifications were transferred between LC-MS/MS runs with the “match between runs” option in which the maximal match time window was set to 0.7 min and the alignment time window set to 20 min. The quantification is based on the “value at maximum” of the extracted ion current. At least two quantitation events were required for a quantifiable protein. Further analysis and filtering of the results was done in Perseus v1.6.10.0. (Tyanova et al. 2016). Comparison of protein group quantities (relative quantification) between different MS runs is based solely on the LFQ’s as calculated by MaxQuant, MaxLFQ algorithm (Cox et al. 2014).

### Data availability

The mass spectrometry proteomics data for the on-bead digestions have been deposited to the ProteomeXchange Consortium via the PRIDE (Vizcaíno et al. 2016) partner repository (https://www.ebi.ac.uk/pride/archive/) with the dataset identifier PXD061382.

### RNA extraction, cDNA synthesis and RT-PCR

Total RNA was extracted from 14d-old Col-0, *tps6-1,* and *tps7-1* seedlings grown vertically on 0.5× MS medium under long-day conditions (16 h light, 110 μmol m^−2^ s^−1^, 22 °C / 8 h dark, 18 °C). Seedlings were harvested at the end of the night, and RNA extraction was performed using the Nucleospin RNA plants purification kit (Macherey-Nagel) according to the manufacturer’s instructions. DNase-treated RNA (1 μg) was reverse transcribed in a total reaction volume of 10 μL using OligoDT and Super Script III Reverse Transcriptase. The resulting cDNA was then diluted to a final volume of 80 µL and used for RT-PCR amplification of *TPS6*, *TPS7*, and *ACTIN* (primers in Supplementary Table S2b). RT-PCR was performed with an annealing temperature of 55 °C for 30 cycles (*TPS6* and *TPS7*) or 26 cycles (*ACTIN*). PCR products were separated on a 1.2% (w/v) agarose gel by electrophoresis at 100 V for 30 minutes.

### Statistical analyses

For statistical analyses of metabolite data, log2 transformed values were used. Student t-test or Welch’s t-test were applied, depending on whether the assumption of equal variance was met. For all other data, the assumption of equal variance and normality was tested and statistical analyses were performed on log10 transformed data whenever that met the assumptions; otherwise, adequate tests were used as indicated in the corresponding figure legend. In all cases a significance level (a) of 0.05 was used. Statistical analyses were performed using GraphPad Prism version 10.4.1 for Windows (GraphPad Software, La Jolla, CA, USA).

## References

Avidan O, Martins MCM, Feil R, Lohse M, Giorgi FM, Schlereth A, Lunn JE, and Stitt M. Direct and indirect responses of the Arabidopsis transcriptome to an induced increase in trehalose 6-phosphate. Plant Physiol. 2024:196(1):409–431. 10.1093/plphys/kiae196

Avonce N, Mendoza-Vargas A, Morett E, and Iturriaga G. Insights on the evolution of trehalose biosynthesis. BMC Evol Biol. 2006:6(1):109. 10.1186/1471-2148-6-109

Baena-González E and Lunn JE. SnRK1 and trehalose 6-phosphate – two ancient pathways converge to regulate plant metabolism and growth. Curr Opin Plant Biol. 2020:55:52–59. 10.1016/j.pbi.2020.01.010

Baena-González E, Rolland F, Thevelein JM, and Sheen J. A central integrator of transcription networks in plant stress and energy signalling. Nature. 2007:448(7156):938–942. 10.1038/nature06069

Belda-Palazón B, Adamo M, Valerio C, Ferreira LJ, Confraria A, Reis-Barata D, Rodrigues A, Meyer C, Rodriguez PL, and Baena-González E. A dual function of SnRK2 kinases in the regulation of SnRK1 and plant growth. Nat Plants. 2020:6(11):1345–1353. 10.1038/s41477-020-00778-w

Belda-Palazón B, Costa M, Beeckman T, Rolland F, and Baena-González E. ABA represses TOR and root meristem activity through nuclear exit of the SnRK1 kinase. Proceedings of the National Academy of Sciences. 2022:119(28). 10.1073/pnas.2204862119

Blanford J, Zhai Z, Baer MD, Guo G, Liu H, Liu Q, Raugei S, and Shanklin J. Molecular mechanism of trehalose 6-phosphate inhibition of the plant metabolic sensor kinase SnRK1. Sci Adv. 2024:10(20). 10.1126/sciadv.adn0895

Broeckx T, Hulsmans S, and Rolland F. The plant energy sensor: evolutionary conservation and divergence of SnRK1 structure, regulation, and function. J Exp Bot. 2016:67(22):6215– 6252. 10.1093/jxb/erw416

Cabib E and Leloir LF. THE BIOSYNTHESIS OF TREHALOSE PHOSPHATE. Journal of Biological Chemistry. 1958:231(1):259–275. 10.1016/S0021-9258(19)77303-7

Carillo P, Feil R, Gibon Y, Satoh-Nagasawa N, Jackson D, Bläsing OE, Stitt M, and Lunn JE. A fluorometric assay for trehalose in the picomole range. Plant Methods. 2013:9(1). 10.1186/1746-4811-9-21

Chantranupong L, Wolfson RL, and Sabatini DM. Nutrient-Sensing Mechanisms across Evolution. Cell. 2015:161(1):67–83. 10.1016/j.cell.2015.02.041

Cox J, Hein MY, Luber CA, Paron I, Nagaraj N, and Mann M. Accurate Proteome-wide Label-free Quantification by Delayed Normalization and Maximal Peptide Ratio Extraction, Termed MaxLFQ. Molecular & Cellular Proteomics. 2014:13(9):2513–2526. 10.1074/mcp.M113.031591

Cox J and Mann M. MaxQuant enables high peptide identification rates, individualized p.p.b.-range mass accuracies and proteome-wide protein quantification. Nat Biotechnol. 2008:26(12):1367–1372. 10.1038/nbt.1511

Cox J, Neuhauser N, Michalski A, Scheltema RA, Olsen J V., and Mann M. Andromeda: A Peptide Search Engine Integrated into the MaxQuant Environment. J Proteome Res. 2011:10(4):1794–1805. 10.1021/pr101065j

Crozet P, Margalha L, Butowt R, Fernandes N, Elias CA, Orosa B, Tomanov K, Teige M, Bachmair A, Sadanandom A, et al. SUMOylation represses SnRK1 signaling in Arabidopsis. The Plant Journal. 2016:85(1):120–133. 10.1111/tpj.13096

Cuadros-Inostroza Á, Caldana C, Redestig H, Kusano M, Lisec J, Peña-Cortés H, Willmitzer L, and Hannah MA. TargetSearch - a Bioconductor package for the efficient preprocessing of GC-MS metabolite profiling data. BMC Bioinformatics. 2009:10. 10.1186/1471-2105-10-428

Delorge I, Figueroa CM, Feil R, Lunn JE, and Van Dijck P. Trehalose-6-phosphate synthase 1 is not the only active TPS in Arabidopsis thaliana. Biochemical Journal. 2015:466:283–290. 10.1042/BJ20141322

Fichtner F and Lunn JE. The Role of Trehalose 6-Phosphate (Tre6P) in Plant Metabolism and Development. Annu Rev Plant Biol. 2021:72(1):737–760. 10.1146/annurev-arplant-050718-095929

Fichtner F, Olas JJ, Feil R, Watanabe M, Krause U, Hoefgen R, Stitt M, and Lunn JE. Functional features of TREHALOSE-6-PHOSPHATE SYNTHASE1, an essential enzyme in Arabidopsis. Plant Cell. 2020:32(6):1949–1972. 10.1105/tpc.19.00837

Figueroa CM, Feil R, Ishihara H, Watanabe M, Kölling K, Krause U, Höhne M, Encke B, Plaxton WC, Zeeman SC, et al. Trehalose 6-phosphate coordinates organic and amino acid metabolism with carbon availability. The Plant Journal. 2016:85(3):410–423. 10.1111/tpj.13114

Gutierrez-Beltran E and Crespo JL. Compartmentalization, a key mechanism controlling the multitasking role of the SnRK1 complex. J Exp Bot. 2022:73(20):7055–7067. 10.1093/jxb/erac315

Hendriks JHM, Kolbe A, Gibon Y, Stitt M, and Geigenberger P. ADP-Glucose Pyrophosphorylase Is Activated by Posttranslational Redox-Modification in Response to Light and to Sugars in Leaves of Arabidopsis and Other Plant Species. Plant Physiol. 2003:133(2):838–849. 10.1104/pp.103.024513

Hennion N, Durand M, Vriet C, Doidy J, Maurousset L, Lemoine R, and Pourtau N. Sugars en route to the roots. Transport, metabolism and storage within plant roots and towards microorganisms of the rhizosphere. Physiol Plant. 2019:165(1):44–57. 10.1111/ppl.12751

Henry C, Bledsoe SW, Siekman A, Kollman A, Waters BM, Feil R, Stitt M, and Lagrimini LM. The trehalose pathway in maize: Conservation and gene regulation in response to the diurnal cycle and extended darkness. J Exp Bot. 2014:65(20). 10.1093/jxb/eru335

Hughes CS, Moggridge S, Müller T, Sorensen PH, Morin GB, and Krijgsveld J. Single-pot, solid-phase-enhanced sample preparation for proteomics experiments. Nat Protoc. 2019:14(1):68–85. 10.1038/s41596-018-0082-x

Hulsmans S, Rodriguez M, De Coninck B, and Rolland F. The SnRK1 Energy Sensor in Plant Biotic Interactions. Trends Plant Sci. 2016:21(8):648–661. 10.1016/j.tplants.2016.04.008

Ishihara H, Alseekh S, Feil R, Perera P, George GM, Niedźwiecki P, Arrivault S, Zeeman SC, Fernie AR, Lunn JE, et al. Rising rates of starch degradation during daytime and trehalose 6-phosphate optimize carbon availability. Plant Physiol. 2022:189(4):1976–2000. 10.1093/plphys/kiac162

James DE, Stöckli J, and Birnbaum MJ. The aetiology and molecular landscape of insulin resistance. Nat Rev Mol Cell Biol. 2021:22(11). 10.1038/s41580-021-00390-6

Van Leene J, Eeckhout D, Gadeyne A, Matthijs C, Han C, De Winne N, Persiau G, Van De Slijke E, Persyn F, Mertens T, et al. Mapping of the plant SnRK1 kinase signalling network reveals a key regulatory role for the class II T6P synthase-like proteins. Nat Plants. 2022:8(11):1245–1261. 10.1038/s41477-022-01269-w

Leverve X, Batandier C, and Fontaine E. Choosing the right substrate. Novartis Found Symp. 2007:280:108–21; discussion 121-7, 160–4.

Lisec J, Schauer N, Kopka J, Willmitzer L, and Fernie AR. Gas chromatography mass spectrometry-based metabolite profiling in plants. Nat Protoc. 2006:1(1). 10.1038/nprot.2006.59

Lunn JE. Gene families and evolution of trehalose metabolism in plants. Functional Plant Biology. 2007:34(6):550. 10.1071/FP06315

Lunn JE. Sucrose Metabolism. . In. Encyclopedia of Life Sciences. (Wiley), pp. 1–9. 10.1002/9780470015902.a0021259.pub2

Lunn JE, Delorge I, Figueroa CM, Van Dijck P, and Stitt M. Trehalose metabolism in plants. Plant Journal. 2014:79(4):544–567. 10.1111/tpj.12509

Lunn JE, Feil R, Hendriks JHM, Gibon Y, Morcuende R, Osuna D, Scheible WR, Carillo P, Hajirezaei MR, and Stitt M. Sugar-induced increases in trehalose 6-phosphate are correlated with redox activation of ADPglucose pyrophosphorylase and higher rates of starch synthesis in Arabidopsis thaliana. Biochemical Journal. 2006:397(1):139–148. 10.1042/BJ20060083

Mair A, Pedrotti L, Wurzinger B, Anrather D, Simeunovic A, Weiste C, Valerio C, Dietrich K, Kirchler T, Nägele T, et al. SnRK1-triggered switch of bZIP63 dimerization mediates the low-energy response in plants. Elife. 2015:4(AUGUST2015). 10.7554/eLife.05828

Margalha L, Confraria A, and Baena-González E. SnRK1 and TOR: Modulating growth–defense trade-offs in plant stress responses. J Exp Bot. 2019:70(8):2261–2274. 10.1093/jxb/erz066

Margalha L, Elias A, Belda-Palazón B, Peixoto B, Confraria A, and Baena-González E. HOS1 promotes plant tolerance to low-energy stress via the SnRK1 protein kinase. Plant Journal. 2023:115(3). 10.1111/tpj.16250

Martínez-Barajas E, Delatte T, Schluepmann H, de Jong GJ, Somsen GW, Nunes C, Primavesi LF, Coello P, Mitchell RAC, and Paul MJ. Wheat Grain Development Is Characterized by Remarkable Trehalose 6-Phosphate Accumulation Pregrain Filling: Tissue Distribution and Relationship to SNF1-Related Protein Kinase1 Activity. Plant Physiol. 2011:156(1):373–381. 10.1104/pp.111.174524

Martins MCM, Hejazi M, Fettke J, Steup M, Feil R, Krause U, Arrivault S, Vosloh D, Figueroa CM, Ivakov A, et al. Feedback inhibition of starch degradation in Arabidopsis leaves mediated by trehalose 6-phosphate. Plant Physiol. 2013:163(3):1142–1163. 10.1104/pp.113.226787

Meijering E, Jacob M, Sarria JCF, Steiner P, Hirling H, and Unser M. Design and Validation of a Tool for Neurite Tracing and Analysis in Fluorescence Microscopy Images. Cytometry Part A. 2004:58(2). 10.1002/cyto.a.20022

Mergner J, Frejno M, List M, Papacek M, Chen X, Chaudhary A, Samaras P, Richter S, Shikata H, Messerer M, et al. Mass-spectrometry-based draft of the Arabidopsis proteome. Nature. 2020:579(7799):409–414. 10.1038/s41586-020-2094-2

Morales-Herrera S, Jourquin J, Coppé F, Lopez-Galvis L, De Smet T, Safi A, Njo M, Griffiths CA, Sidda JD, Mccullagh JSO, et al. Trehalose-6-phosphate signaling regulates lateral root formation in Arabidopsis thaliana. Proceedings of the National Academy of Sciences. 2023:120(40). 10.1073/pnas.2302996120

Muralidhara P, Weiste C, Collani S, Krischke M, Kreisz P, Draken J, Feil R, Mair A, Teige M, Müller MJ, et al. Perturbations in plant energy homeostasis prime lateral root initiation via SnRK1-bZIP63-ARF19 signaling. Proc Natl Acad Sci U S A. 2021:118(37). 10.1073/pnas.2106961118

Nozue K, Covington MF, Duek PD, Lorrain S, Fankhauser C, Harmer SL, and Maloof JN. Rhythmic growth explained by coincidence between internal and external cues. Nature. 2007:448(7151):358–361. 10.1038/nature05946

Nunes C, O’Hara LE, Primavesi LF, Delatte TL, Schluepmann H, Somsen GW, Silva AB, Fevereiro PS, Wingler A, and Paul MJ. The trehalose 6-phosphate/snRK1. signaling pathway primes growth recovery following relief of sink limitation. Plant Physiol. 2013a:162(3). 10.1104/pp.113.220657

Nunes C, Primavesi LF, Patel MK, Martinez-Barajas E, Powers SJ, Sagar R, Fevereiro PS, Davis BG, and Paul MJ. Inhibition of SnRK1 by metabolites: Tissue-dependent effects and cooperative inhibition by glucose 1-phosphate in combination with trehalose 6-phosphate. Plant Physiology and Biochemistry. 2013b:63:89–98. 10.1016/j.plaphy.2012.11.011

Pedrotti L, Weiste C, Nägele T, Wolf E, Lorenzin F, Dietrich K, Mair A, Weckwerth W, Teige M, Baena-González E, et al. Snf1-RELATED KINASE1-controlled C/S1-bZIP signaling activates alternative mitochondrial metabolic pathways to ensure plant survival in extended darkness. Plant Cell. 2018:30(2):495–509. 10.1105/tpc.17.00414

Peixoto B and Baena-González E. Management of plant central metabolism by SnRK1 protein kinases. J Exp Bot. 2022:73(20):7068–7082. 10.1093/jxb/erac261

Peixoto B, Moraes TA, Mengin V, Margalha L, Vicente R, Feil R, Höhne M, Sousa AGG, Lilue J, Stitt M, et al. Impact of the SnRK1 protein kinase on sucrose homeostasis and the transcriptome during the diel cycle. Plant Physiol. 2021:187(3):1357–1373. 10.1093/plphys/kiab350

Ramon M, Dang TVT, Broeckx T, Hulsmans S, Crepin N, Sheen J, and Rolland F. Default Activation and Nuclear Translocation of the Plant Cellular Energy Sensor SnRK1 Regulate Metabolic Stress Responses and Development. Plant Cell. 2019:31(7):1614–1632. 10.1105/tpc.18.00500

Ramon M, De Smet I, Vandesteene L, Naudts M, Leyman B, Van Dijck P, Rolland F, Beeckman T, and Thevelein JM. Extensive expression regulation and lack of heterologous enzymatic activity of the Class II trehalose metabolism proteins from Arabidopsis thaliana. Plant Cell Environ. 2009:32(8):1015–1032. 10.1111/j.1365-3040.2009.01985.x

Rappsilber J, Mann M, and Ishihama Y. Protocol for micro-purification, enrichment, pre-fractionation and storage of peptides for proteomics using StageTips. Nat Protoc. 2007:2(8):1896–1906. 10.1038/nprot.2007.261

Roessner U, Luedemann A, Brust D, Fiehn O, Linke T, Willmitzer L, and Fernie AR. Metabolic profiling allows comprehensive phenotyping of genetically or environmentally modified plant systems. Plant Cell. 2001:13(1). 10.1105/tpc.13.1.11

Sanagi M, Aoyama S, Kubo A, Lu Y, Sato Y, Ito S, Abe M, Mitsuda N, Ohme-Takagi M, Kiba T, et al. Low nitrogen conditions accelerate flowering by modulating the phosphorylation state of FLOWERING BHLH 4 in *Arabidopsis*. Proceedings of the National Academy of Sciences. 2021:118(19). 10.1073/pnas.2022942118

Shen W, Reyes MI, and Hanley-Bowdoin L. Arabidopsis Protein Kinases GRIK1 and GRIK2 Specifically Activate SnRK1 by Phosphorylating Its Activation Loop. Plant Physiol. 2009:150(2):996–1005. 10.1104/pp.108.132787

Sulpice R, Flis A, Ivakov AA, Apelt F, Krohn N, Encke B, Abel C, Feil R, Lunn JE, and Stitt M. Arabidopsis coordinates the diurnal regulation of carbon allocation and growth across a wide range of photoperiods. Mol Plant. 2014:7(1):137–155. 10.1093/mp/sst127

Tyanova S, Temu T, Sinitcyn P, Carlson A, Hein MY, Geiger T, Mann M, and Cox J. The Perseus computational platform for comprehensive analysis of (prote)omics data. Nat Methods. 2016:13(9):731–740. 10.1038/nmeth.3901

Vandesteene L, Ramon M, Le Roy K, Van Dijck P, and Rolland F. A Single Active Trehalose-6-P Synthase (TPS) and a Family of Putative Regulatory TPS-Like Proteins in Arabidopsis. Mol Plant. 2010:3(2):406–419. 10.1093/mp/ssp114

Vizcaíno JA, Csordas A, del-Toro N, Dianes JA, Griss J, Lavidas I, Mayer G, Perez-Riverol Y, Reisinger F, Ternent T, et al. 2016 update of the PRIDE database and its related tools. Nucleic Acids Res. 2016:44(D1):D447–D456. 10.1093/nar/gkv1145

Vogel G, Fiehn O, Jean-Richard-dit-Bressel L, Boller T, Wiemken A, Aeschbacher RA, and Wingler A. Trehalose metabolism in Arabidopsis: occurrence of trehalose and molecular cloning and characterization of trehalose-6-phosphate synthase homologues. J Exp Bot. 2001:52(362):1817–1826. 10.1093/jexbot/52.362.1817

Wahl V, Ponnu J, Schlereth A, Arrivault S, Langenecker T, Franke A, Feil R, Lunn JE, Stitt M, and Schmid M. Regulation of flowering by trehalose-6-phosphate signaling in Arabidopsis thaliana. Science (1979). 2013:339(6120):704–707. 10.1126/science.1230406

Wingler A, Delatte TL, O’Hara LE, Primavesi LF, Jhurreea D, Paul MJ, and Schluepmann H. Trehalose 6-Phosphate Is Required for the Onset of Leaf Senescence Associated with High Carbon Availability. Plant Physiol. 2012:158(3):1241–1251. 10.1104/pp.111.191908

Xiang C, Han P, Lutziger I, Wang K, and Oliver DJ. A mini binary vector series for plant transformation. Plant Mol Biol. 1999:40(4). 10.1023/A:1006201910593

Yadav UP, Ivakov A, Feil R, Duan GY, Walther D, Giavalisco P, Piques M, Carillo P, Hubberten HM, Stitt M, et al. The sucrose-trehalose 6-phosphate (Tre6P) nexus: Specificity and mechanisms of sucrose signalling by Tre6P. J Exp Bot. 2014:65(4):1051–1068. 10.1093/jxb/ert457

Yazdanbakhsh N, Sulpice R, Graf A, Stitt M, and Fisahn J. Circadian control of root elongation and C partitioning in *Arabidopsis thaliana*. Plant Cell Environ. 2011:34(6):877–894. 10.1111/j.1365-3040.2011.02286.x

Zacharaki V, Ponnu J, Crepin N, Langenecker T, Hagmann J, Skorzinski N, Musialak-Lange M, Wahl V, Rolland F, and Schmid M. Impaired KIN10 function restores developmental defects in the *Arabidopsis trehalose 6-phosphate synthase1 (tps1)* mutant. New Phytologist. 2022:235(1):220–233. 10.1111/nph.18104

Zhai Z, Keereetaweep J, Liu H, Feil R, Lunn JE, and Shanklin J. Trehalose 6-phosphate positively regulates fatty acid synthesis by stabilizing wrinkled1. Plant Cell. 2018:30(10):2616–2627. 10.1105/tpc.18.00521

Zhang Y, Primavesi LF, Jhurreea D, Andralojc PJ, Mitchell RAC, Powers SJ, Schluepmann H, Delatte T, Wingler A, and Paul MJ. Inhibition of SNF1-Related Protein Kinase1 Activity and Regulation of Metabolic Pathways by Trehalose-6-Phosphate. Plant Physiol. 2009:149(4):1860–1871. 10.1104/pp.108.133934

Zhang ZP, Deng Y, Song X, and Miao M. Trehalose-6-phosphate and SNF1-related protein kinase 1 are involved in the first-fruit inhibition of cucumber. J Plant Physiol. 2015:177. 10.1016/j.jplph.2014.09.009

