## Supplementary Figures for "TPS Proteins coordinate plant growth with sugar availability via the SnRK1 Kinase"

### Supplementary Figure 1

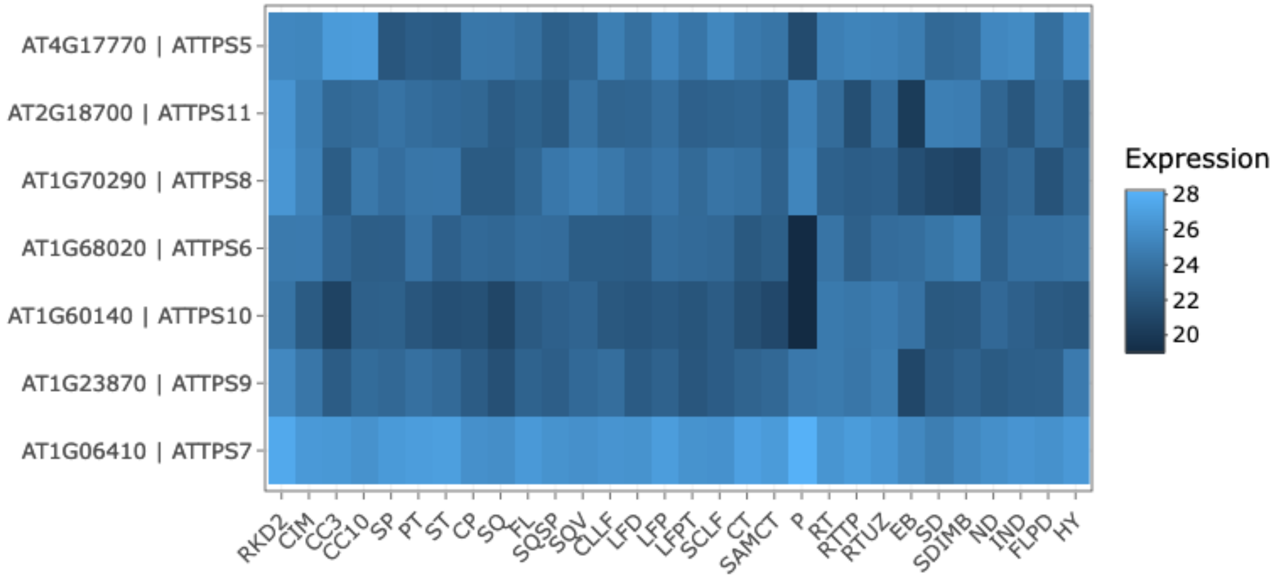


**Supplementary Figure 1. Expression atlas of class II TPS proteins in Arabidopsis.** Relative protein abundance in different tissues based on data from Mergner et al. (2020) and visualized with ATHENA proteomics (https://athena.proteomics.wzw.tum.de/master_arabidopsisshiny/). *RKD2*, egg-cell like callus; *CIM*, callus; *CC3*, root cell culture early; *CC10*, root cell culture late; *SP*, sepal (flower stage 15); *PT*, petal (flower stage 15); *ST*, stamen (flower stage 15); *CP*, carpel (flower stage 15); *SQ*, silique (stage 3); *FL*, flower (stage 15); *SQSP*, silique septum; *SQV*, silique valves; *CLLF*, 1st cauline leaf; *LFD*, rosette leaf 7, distal part; *LFP*, rosette leaf 7, proximal part; *LFPT*, rosette leaf 7, petiole; *SCLF*, sensecent leaf; *CT*, cotyledons; *SAMCT*, shoot apical meristem, cotyledons and first leaves; *P*, pollen; *RT*, root; *RTP*, root tip; *RTUZ*, root upper zone; *EB*, seed (embryo stage 10); *SD*, seed (mature, dry); *SDIMB*, seed (mature, imbibed); *ND*, stem, 1st node; *IND*, stem, 2nd internode; *FLPD*, flower pedicle (flower stage 15); *HY*, hypocotyl.

### Supplementary Figure 2

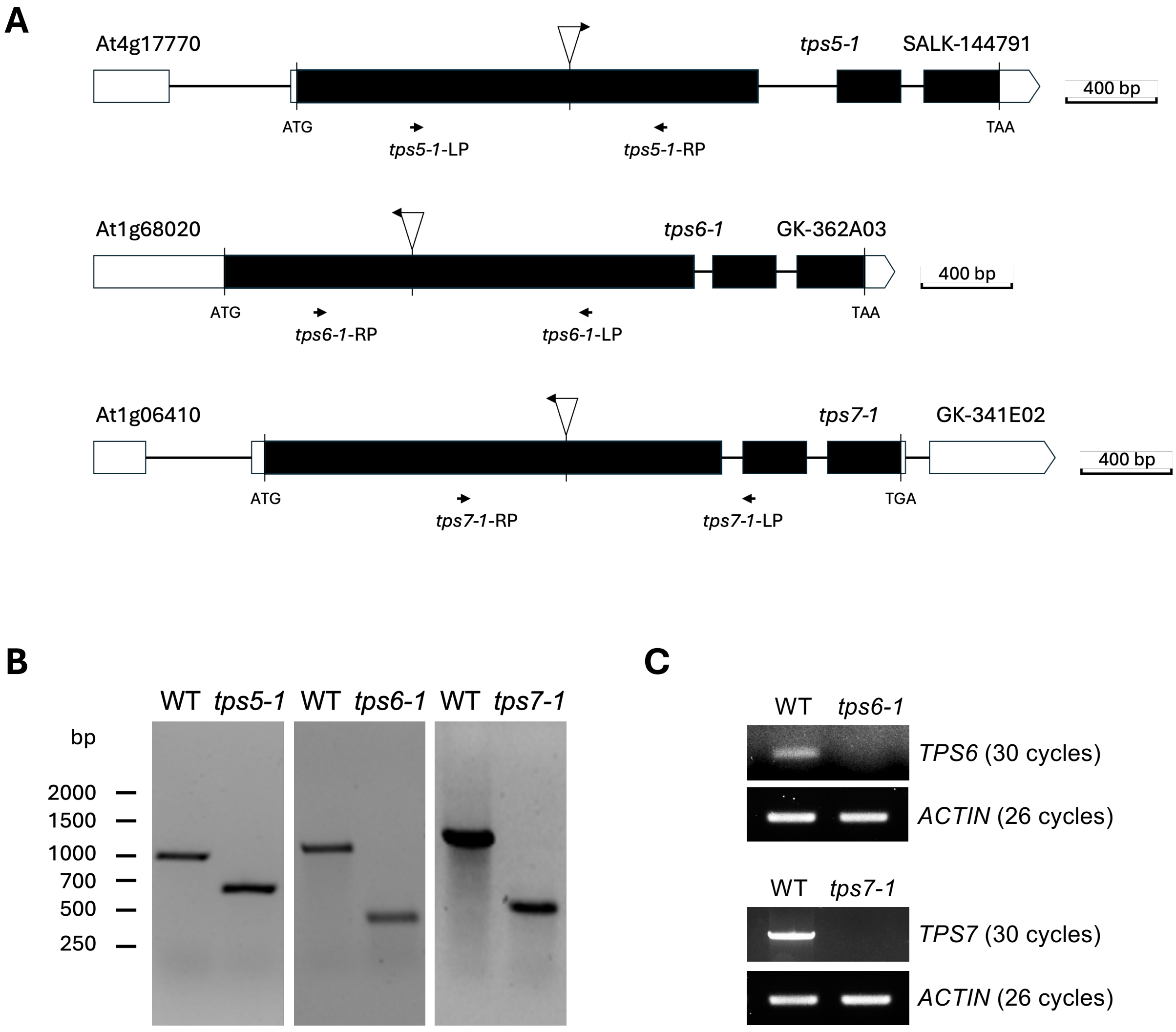


**Supplementary Figure 2. Generation of *tps* mutants.** (**A**) Schematic representation of the insertion sites of the *tps5*, *tps6*, and *tps7* T-DNA mutants used in this study. The ATG start codon and TAA/TGA stop codons are indicated. Exons are shown as black boxes, introns as black lines, and T-DNA insertion sites as triangles above the corresponding gene diagrams. Scale bar: 400 bp. (**B**) Confirmation of *tps* mutant identities by genotyping using T-DNA left border and gene-specific primers (LP and RP, indicated in **A**). (**C**) RT-PCR analyses of the *TPS6* (upper panel) and *TPS7* (lower panel) transcripts in *tps6-1* and *tps7-1* mutants, respectively. Amplification of *ACTIN* mRNA was used as a control.

### Supplementary Figure 3

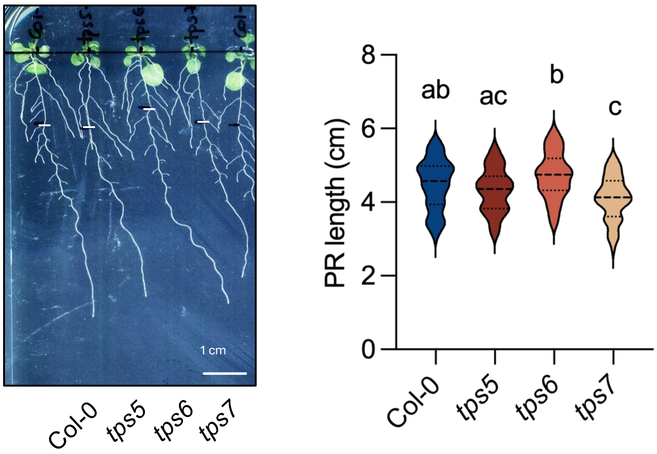


**Supplementary Figure 3. Phenotype of single *tps5*, *tps6*, and *tps7* mutants.** Left, representative images of 14d-old Col-0, *tps5*, *tps6* and *tps7* seedlings grown vertically on 0.5x MS medium for 7 d and transferred to 0.5x MS medium for 7 d. White marks indicate root length at transfer. Scale bar 1 cm. Right, quantification of primary root (PR) length, measured from the white mark to the root tip ($n$=3 independent experiments, 48-64 seedlings per genotype). In the violin plot, dotted lines represent the quartiles and the dashed line, the median. Different letters indicate statistically significant differences between genotypes ($p$<0.05, one-way ANOVA with Tukey HSD test).

### Supplementary Figure 4

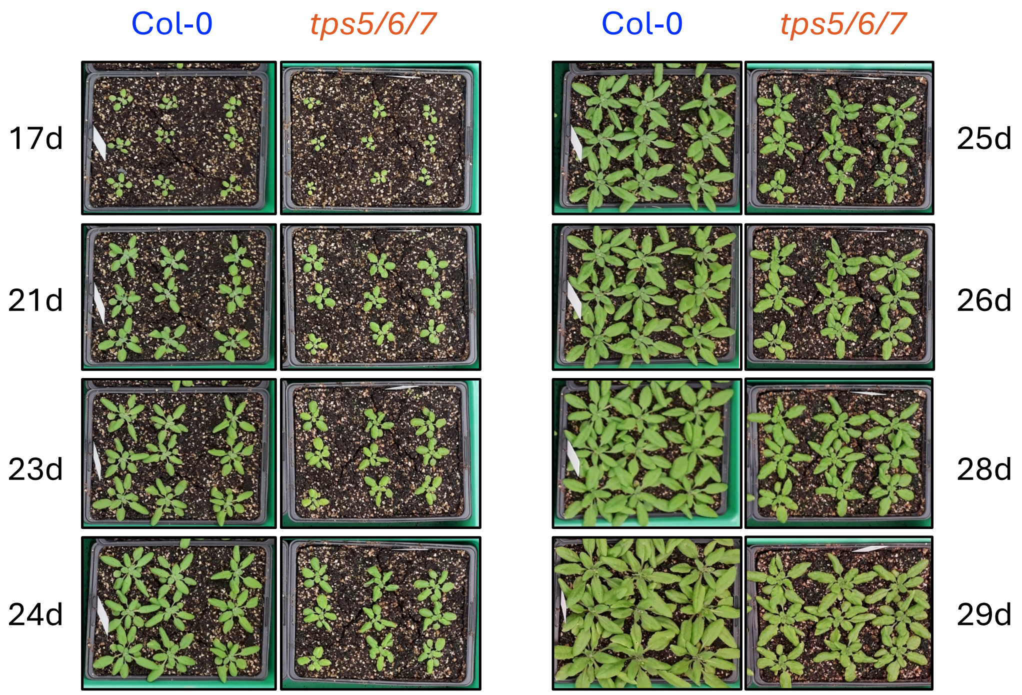


**Supplementary Figure 4. Rosette phenotype of Col-0 and *tps5/6/7* plants.** Representative images of rosettes from Col-0 (left) and *tps5/6/7* (right) plants grown on soil under long-day conditions (16 h light, 110 µmol $m^{-2}$ $s^{-1}$, 22 °C / 8 h dark, 18 °C). Images were captured over 12 days, from day 17 to day 29 after sowing

### Supplementary Figure 5

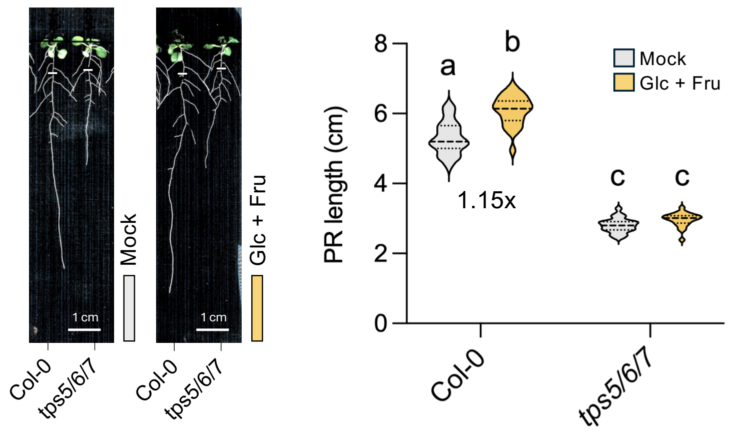


**Supplementary Figure 5. Impact of glucose and fructose on the *tps5/6/7* PR growth phenotype.** Left, representative images of 14d-old Col-0 and *tps5/6/7* seedlings grown vertically on 0.5x MS medium for 7 d and transferred to 0.5x MS medium supplemented (right) or not (left) with 30 mM Glucose + 30 mM fructose for 7 d. White marks indicate root length at transfer. Scale bar 1 cm. Right, quantification of PR length, measured from the white mark to the root tip (n=1 independent experiment, 22-24 seedlings per genotype per condition). In the violin plot, dotted lines represent the quartiles and the dashed line, the median. Different letters indicate statistically significant differences between genotype and condition combinations ($p$<0.05, one-way ANOVA with Tukey HSD test).

### Supplementary Figure 6

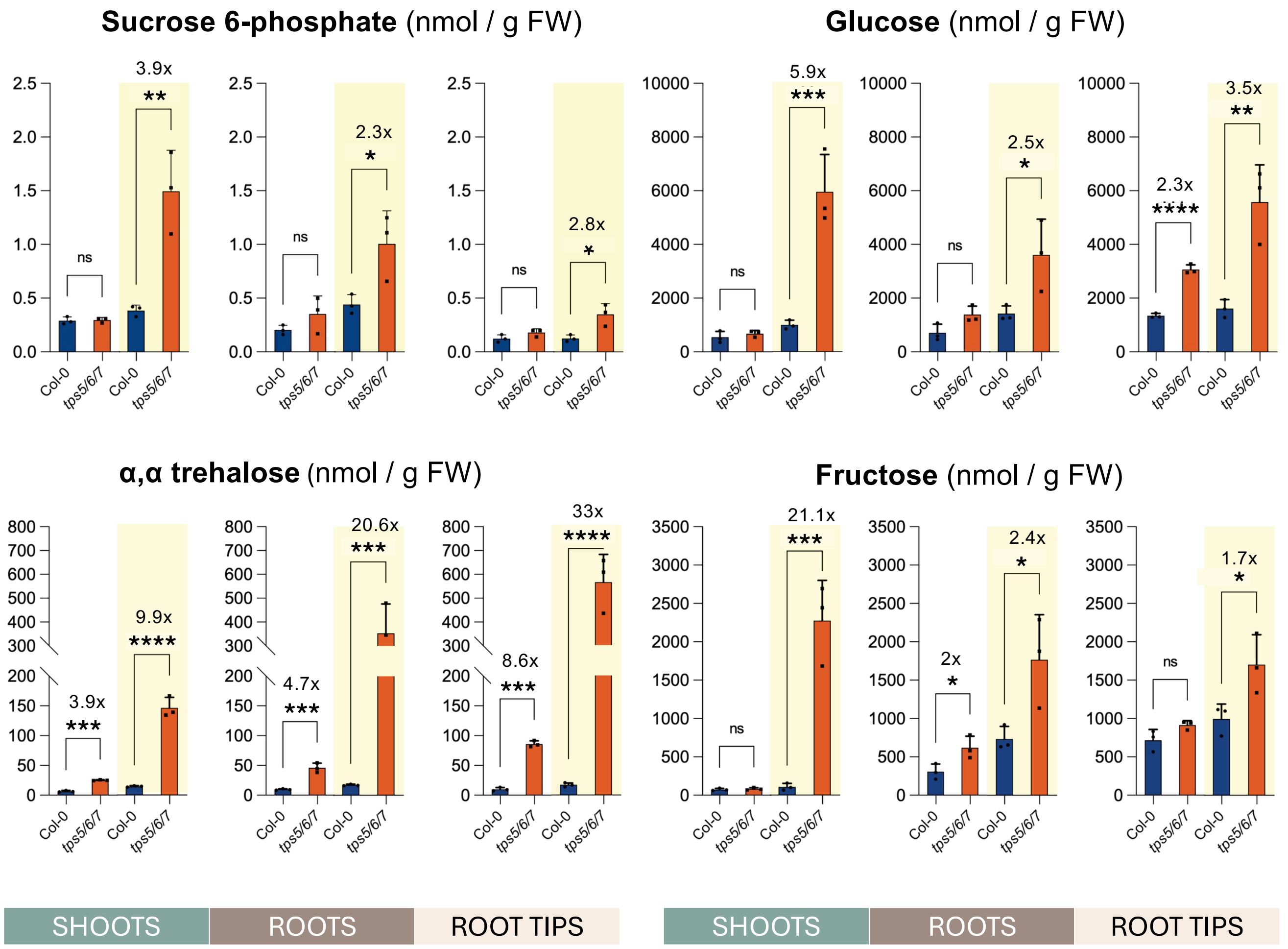


**Supplementary Figure 6. Impact of TPS5/6/7 depletion on sugars related to sucrose and trehalose metabolism.** Levels of sucrose 6-phosphate, glucose, fructose, and α,α trehalose in the indicated tissues of 14d-old Col-0 and *tps5/6/7* seedlings. Seedlings were grown vertically on 0.5x MS medium under control light conditions (110 μmol $m^{-2}$ $s^{-1}$) for 7 d, transferred to fresh 0.5x MS plates and grown for 7 d under control light (CL; left, white) or high light conditions (HL; 200 μmol $m^{-2}$ $s^{-1}$; right, yellow). Graphs show the mean of 3 independent experiments (each consisting of 200-250 seedlings per genotype per condition, error bars, SD). Asterisks indicate statistically significant differences between genotypes within each tissue and condition ($p$<0.05, two-tailed unpaired Student t-test).

### Supplementary Figure 7

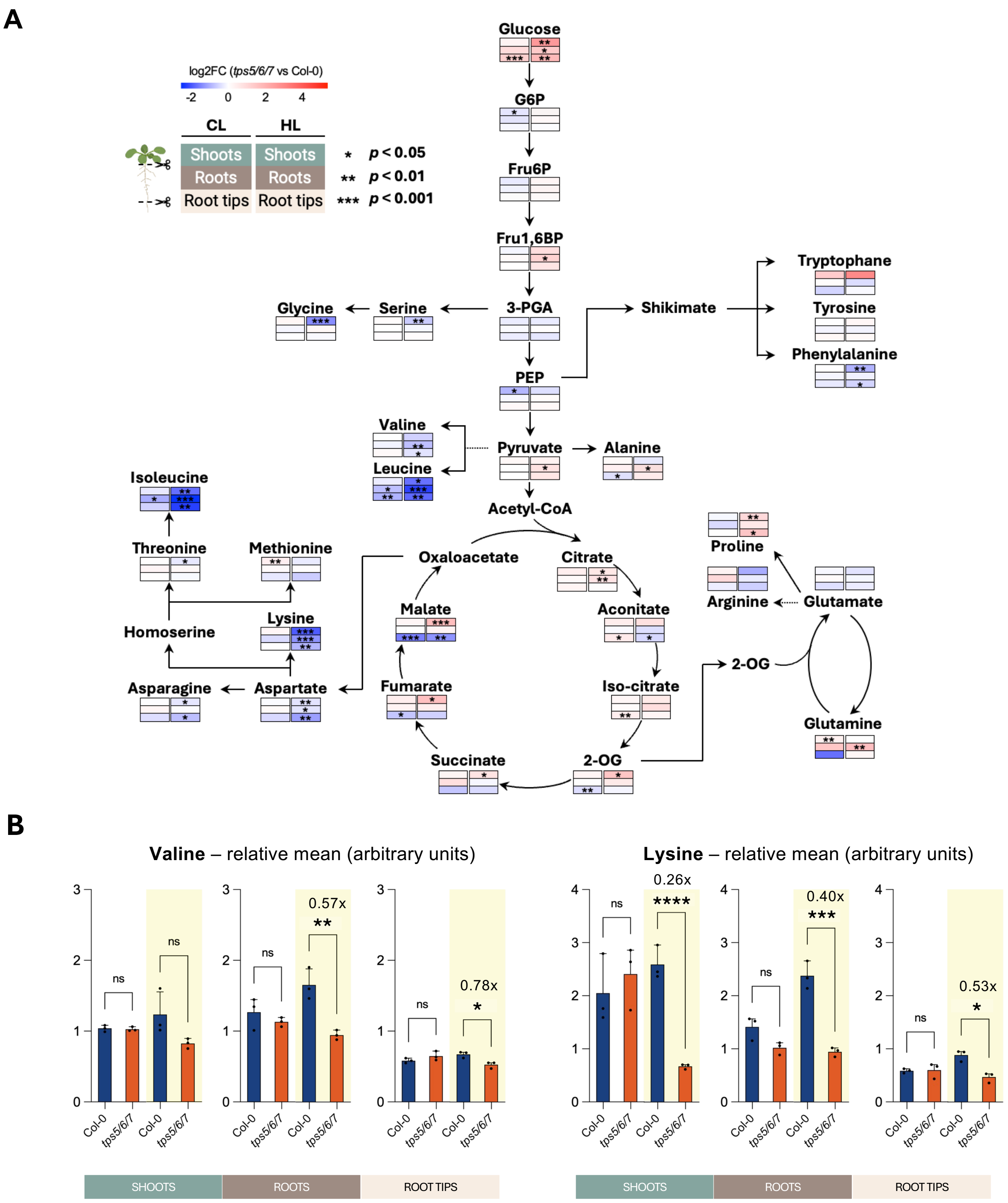


**Supplementary Figure 7. Impact of TPS5/6/7 depletion on primary metabolism.** (**A**) Ratios of primary metabolites in the indicated tissues of 14d-old Col-0 and *tps5/6/7* seedlings. Seedlings were grown vertically on 0.5x MS medium under control light conditions (110 µmol $m^{-2} s^{-1}$) for 7 d, transferred to fresh 0.5x MS plates and grown for 7 d under control light (CL; left) or high light conditions (HL; 200 µmol $m^{-2} s^{-1}$; right). Colors indicate the log2 ratios of the mean values (*tps5/6/7 vs.* Col-0; *n*=3). Asterisks indicate statistically significant changes in metabolite levels between genotypes within each tissue and light condition (**p*<0.05, ***p*<0.01, ****p*<0.001; two-tailed unpaired Student and Welch's t-tests). (**B**) Levels of Lysine and Valine in the indicated tissues of 14d-old Col-0 and *tps5/6/7* seedlings grown as in (**A**) under contrasting light intensities (CL, white; HL, yellow). Graphs show the mean of 3 independent experiments (each consisting of 200-250 seedlings per genotype per condition, error bars, SD). Asterisks indicate statistically significant differences between genotypes within each tissue and condition (*p*<0.05, two-tailed unpaired Student and Welch's t-tests).

### Supplementary Figure 8

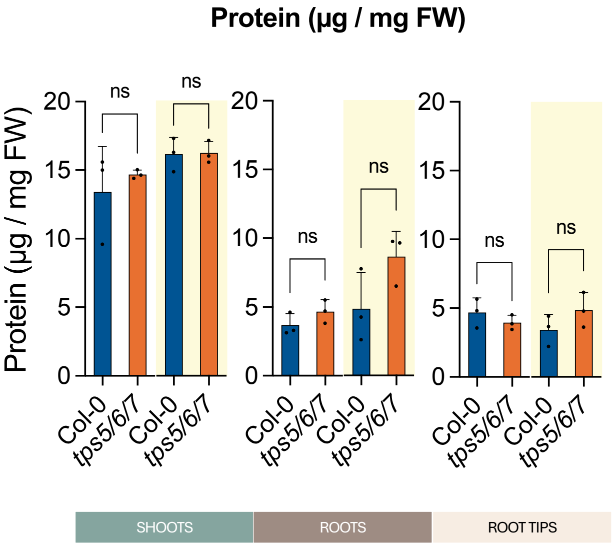


**Supplementary Figure 8. Impact of TPS5/6/7 depletion on total protein content.** Total protein levels in the indicated tissues of 14d-old Col-0 and *tps5/6/7* seedlings. Seedlings were grown vertically on 0.5x MS medium under control light conditions (110 µmol $m^{-2} s^{-1}$) for 7 d, transferred to fresh 0.5x MS plates and grown for an additional 7 d under control light (CL; left, white) or high light conditions (HL; 200 µmol $m^{-2} s^{-1}$; right, yellow). Samples were harvested at the end of the night. Graphs show the mean of 3 independent experiments (each consisting of 200-250 seedlings per genotype per condition, error bars, SD). Asterisks indicate statistically significant differences between genotypes within each tissue and condition ($p$<0.05, two-tailed unpaired Student and Welch t-tests).

### Supplementary Figure 9

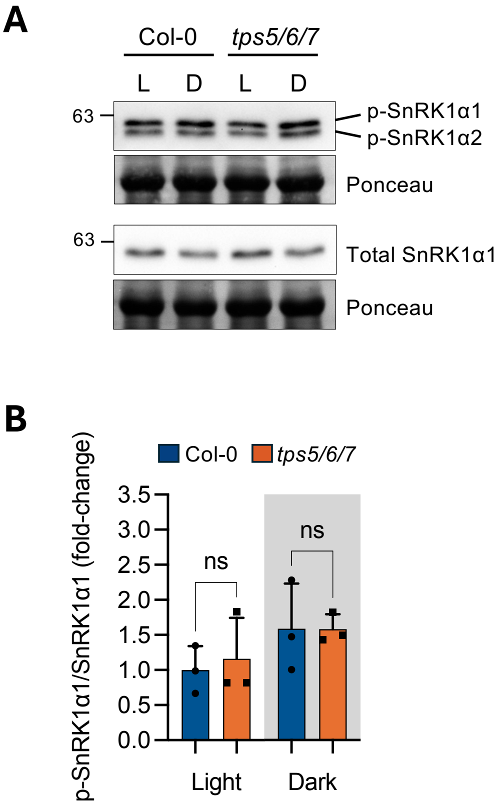


**Supplementary Figure 9. Impact of TPS5/6/7 depletion on SnRK1α1 T-loop phosphorylation. (A)** Representative immunoblots showing SnRK1α T-loop phosphorylation (p-SnRK1α1 and p-SnRK1α2) and total SnRK1α1 levels detected, respectively, with anti-phosphoAMPK (p-AMPK) and anti-SnRK1α1 antibodies. Proteins were extracted from 14d-old Col-0 and *tps5/6/7* seedlings, either maintained in the light (L) or transferred to darkness (D) at ZT2 for 4 h and harvested at ZT6. **(B)** Quantification of SnRK1α1 T-loop phosphorylation, presented as the p-SnRK1α1/SnRK1α1 ratio. Bars show the mean of 3 independent experiments (error bars, SD). Asterisks indicate significant differences between genotypes ($p$<0.05; two tailed unpaired Student t-test).

### Supplementary Figure 10

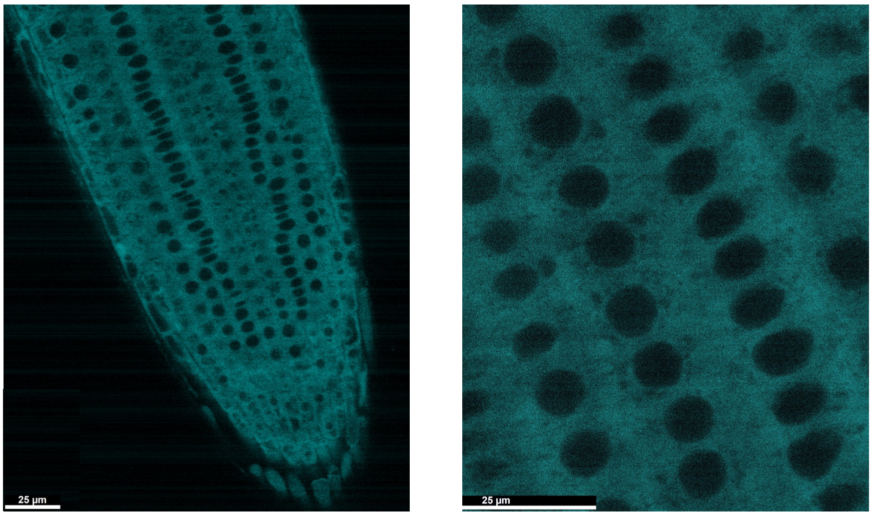


**Supplementary Figure 10. Subcellular localization of TPS7-mCherry.** Representative fluorescence microscopy images of the root meristem of Arabidopsis seedlings stably expressing *TPS7-mCherry* (line C#2). Scale bar: 25 µm.

### Supplementary Figure 11

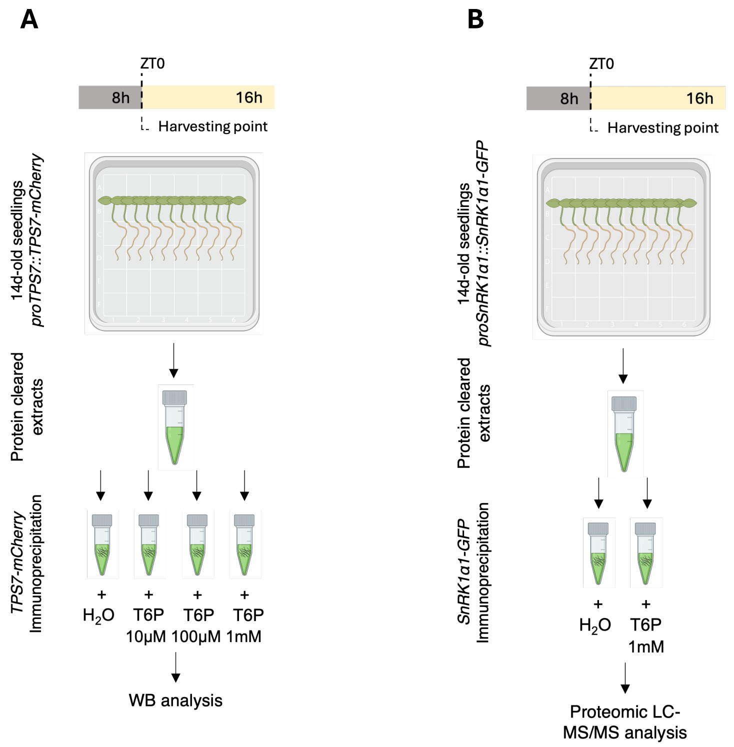


**Supplementary Figure 11. Experimental set-up for assessing the TPS7-SnRK1α1 interaction.** Cleared protein extracts from 14d-old *proTPS7::TPS7-mCherry* (**A**) or *proSnRK1α1::SnRK1α1-GFP* (**B**) seedlings, harvested at EN, were aliquoted and incubated with ChromoTek RFP- or GFP-Trap® Magnetic Agarose beads, respectively, in the absence ($H_2O$ control) or presence of the indicated concentrations of T6P. (**A**) Immunoprecipitated mCherry-tagged TPS7 and co-immunoprecipitated SnRK1α1 were assessed by immunodetection (Fig. 4A-B). (**B**) Proteins interacting with GFP-tagged SnRK1α1 were identified by MS/MS analyses (Fig. 4C).
